# Dorsomedial striatal GABA dynamics organize palatable reward consumption and are reshaped by GLP-1 receptor agonism

**DOI:** 10.64898/2026.09.03.749293

**Authors:** Aryanna Nichelle Copling, Oyku Dinckol, Noah Harris Wenger, Bhumiben Pradipkumar Patel, Ezechiel Muanza N’Kongolo Lukusa, Munir Gunes Kutlu

**Author notes:** These authors contributed equally (Aryanna Nichelle Copling, Oyku Dinckol). **Corresponding Author Munir Gunes Kutlu, PhD** Assistant Professor Center for Substance Abuse Research (CSAR) Department of Neural Sciences Temple University Lewis Katz School of Medicine 3500 N Broad St, Philadelphia, PA 19140.

## Abstract

Palatability and metabolic state strongly shape food consumption, making it important to understand the neural mechanisms that integrate these influences. Glucagon-like peptide-1 (GLP-1) receptor agonists are potent modulators of food intake and increasingly used therapeutic options, yet the circuit mechanisms underlying their effects remain unclear. Dorsal striatal inhibitory circuits contribute to reward-guided behavior and feeding, but how they encode ongoing consumption or are altered by GLP-1 receptor agonism is unknown. Using fiber photometry in mice, we found that dorsomedial striatal (DMS) GABA signals decreased at consumption onset, scaled with palatability, predicted licking, and were enhanced by food deprivation. GLP-1 receptor agonist semaglutide reduced intake, disrupted coupling between DMS GABA and licking, enhanced rebound signals preceding pauses, and increased DMS ensemble synchrony. Optogenetic excitation of DMS GABAergic interneurons reproduced key semaglutide-induced changes in consumption structure. These findings suggest that DMS GABA dynamics, as a state-dependent regulator of palatable consumption, are reshaped by GLP-1 receptor agonism.

## Main Text

Many brain regions contribute to various aspects of food intake, including palatability, motivation, food choice, and satiety^1,2^. Among these is the striatum, comprising both the ventral and dorsal components, which plays a central role in integrating motivational signals with motor output to guide goal-directed behavior^3–6^. The striatum is a GABA-dominated circuit, in which the vast majority of neurons release γ-aminobutyric acid (GABA), making inhibitory signaling the primary mode of information processing^7–9^. The striatum includes five principal neuronal cell types, including two classes of GABAergic projection neurons, known as medium spiny neurons (MSNs), and three major interneuron populations.^10–12^ These interneuron populations include fast-spiking interneurons (FSIs), which provide strong feedforward inhibition onto MSNs; low-threshold spiking interneurons (LTSIs), which contribute to sustained inhibitory tone and modulatory control; and cholinergic interneurons (CINs), which indirectly regulate GABA release through modulation of presynaptic inputs and local circuit excitability^13–17^. Within this microcircuit, GABA release arises from both MSN axon collaterals and interneuron-driven inhibition, generating tightly timed lateral and feedforward inhibition that shapes MSN ensemble activity and output selection^18^. In addition to its intrinsic inhibitory circuitry, the striatum receives inhibitory inputs from local interneurons, MSN axon collaterals^19^, and inputs from other brain regions, such as the cortex^20^ and subcortical structures, including the ventral pallidum^21^, among others^22,23^. Collectively, these properties establish GABA release as one of the principal determinants of striatal circuit dynamics and output.

Building on this circuitry, the dorsal striatum is well positioned to translate reward-related information into consummatory behavior. The dorsomedial striatum (DMS) has been implicated in encoding action–outcome relationships and reward value, including the hedonic properties that drive consumption of palatable solution^24,25^. In contrast, the dorsolateral striatum (DLS) circuitry is thought to contribute to the reinforcement and automatization of consummatory behaviors, such as repeated licking or intake of highly palatable solutions with experience^26^. GABAergic signaling is a major determinant of striatal output, shaping MSN activity and local circuit interactions that influence action selection and reinforcement^6,26–29^. Accordingly, striatal GABAergic transmission may provide a mechanism through which motivational and reward-related information is translated into ongoing feeding behavior^14,30^. However, despite extensive evidence linking dorsal striatal circuits to reward-guided action and consumption, how extracellular GABA dynamics evolve during ongoing intake, whether these signals differ across dorsal striatal subregions, and how they relate to the palatability and temporal structure of consumption remain largely unknown.

In parallel with these striatal mechanisms, metabolic signals critically regulate food intake by modulating neural circuits that encode reward value and motivation^1,2^. Among these, glucagon-like peptide-1 (GLP-1) receptor signaling has emerged as a powerful regulator of feeding behavior, acting both peripherally and centrally to suppress appetite and reduce the rewarding properties of food.^31,32^ GLP-1 receptors (GLP-1R) are expressed in multiple brain regions, including mesolimbic and striatal circuits, where their activation has been shown to decrease food intake, attenuate reward-driven behaviors, and alter neural responses to palatable stimuli^33,34^. Pharmacological activation of GLP-1 receptors, such as with the clinically used agonist semaglutide, robustly reduces consumption of palatable foods, in part by decreasing hedonic value^35^. However, the circuit-level mechanisms through which GLP-1 signaling alters reward encoding and consumption dynamics within the striatum remain incompletely understood.

These region-specific roles suggest that GABAergic transmission in the dorsal striatum links the hedonic evaluation of palatable solutions to the execution and reinforcement of feeding behaviors in rodents. However, how GABAergic dynamics in the dorsal striatum are engaged during ongoing consumption, and how these signals are modulated by changes in reward value, remain poorly understood. Importantly, although GLP-1 receptor activation is known to suppress food intake and decrease motivation for palatable rewards, whether these effects are mediated by modulation of striatal GABAergic signaling, and how such modulation reshapes real-time consumption dynamics, remain unknown. Thus, in the present study, we combined fiber photometry, single-cell calcium imaging, pharmacological manipulation, and optogenetics to define the role of dorsal striatal GABA signaling during palatable reward consumption. We monitored GABA dynamics in DLS and DMS during ongoing intake and tested how the GLP-1 receptor agonist, semaglutide, alters DMS GABA release and neural ensemble structure. Finally, we examined whether optogenetic stimulation of DMS GABA interneurons could recapitulate key behavioral effects of semaglutide, including reductions in lick bout duration.

## Results

### Dorsomedial but not dorsolateral striatal GABA release scales with the palatability of liquids

We first characterized the GABA dynamics in the dorsal striatum (DLS and DMS) during the consumption of water, Ensure, and saccharin. To achieve this, we utilized fiber photometry recordings in awake, behaving mice expressing the genetically encoded GABA sensor iGABA.Sn.FR2 (pGP-AAV5-syn-iGABA.Sn.FR2, Janelia; see **Fig. 1a** and **Extended Data Fig. 1a** for fiber placements) to monitor real-time extracellular GABA transients in dorsal striatal subregions, including the DMS and DLS. GABA signals were recorded across consecutive days, during which animals had free access to water, Ensure, and saccharin (see **Fig.1b,c** for task schematic) under ad libitum feeding conditions, which vary in palatability, with Ensure providing a calorically dense, high-fat/high-sugar solution, saccharin offering non-caloric sweetness, and water serving as a neutral control. As expected, animals showed progressively higher lick counts with increasing palatability of the solution (**Fig. 1d**; Ensure > saccharin > water). GABA release in both dorsal striatal subregions showed an initial decrease during the first 5 licks, independent of solution type (**Fig. 1e,g**). Notably, DMS GABA dynamics scaled with palatability, with Ensure producing the largest decrease, followed by saccharin and then water (**Fig. 1g,h**). In contrast, DLS GABA decreases did not differ across solutions (**Fig. 1e,f**), indicating a DMS-specific palatability-dependent modulation of GABA signaling. Given the absence of palatability-dependent modulation in DLS GABA release, subsequent analyses were restricted to DMS.

**Figure 1:**
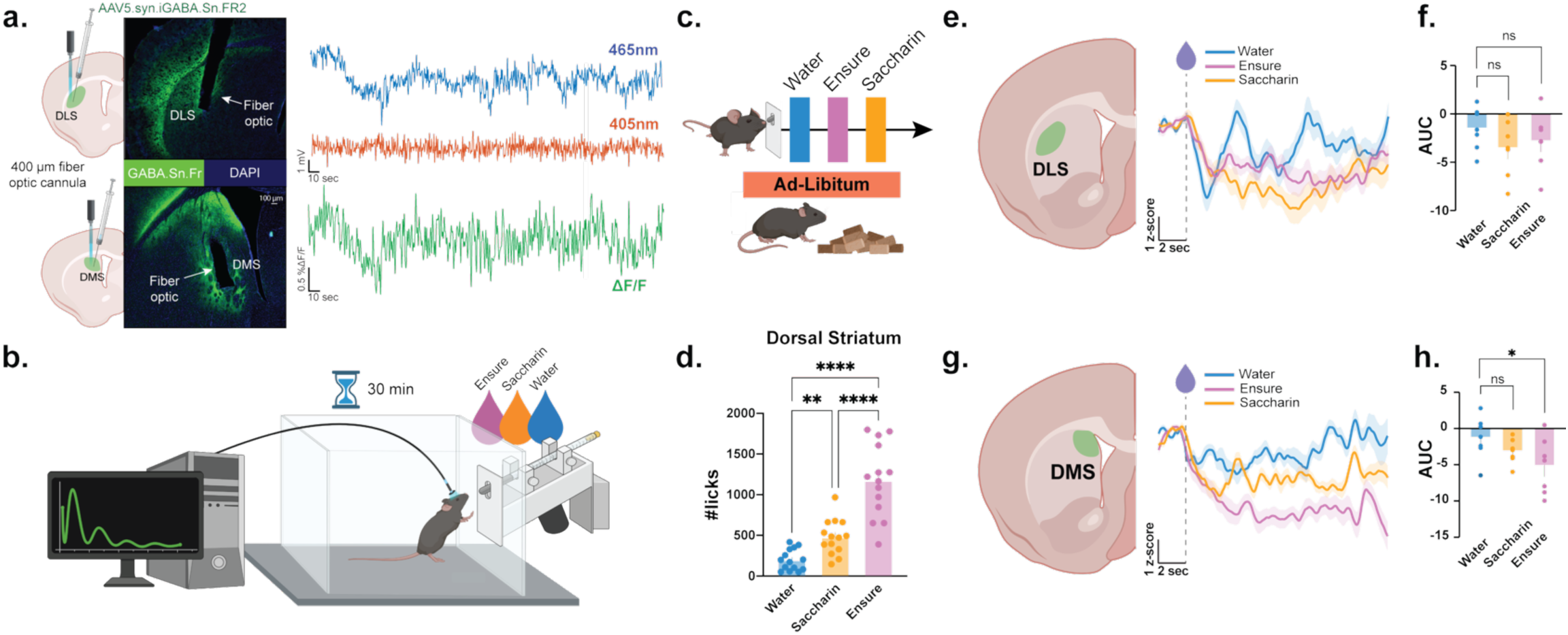
Dorsomedial but not dorsolateral striatal GABA release scales with the palatability of liquids. **(a)** Experimental schematic illustrating unilateral injection of AAV5.syn.iGABA.Sn.FR2 into either the dorsolateral striatum (DLS; *n =* 7, 4 males and 3 females) or dorsomedial striatum (DMS; *n =* 7, 4 males and 3 females), followed by implantation of a 400µm core fiber optic cannula for fiber photometry recordings of extracellular GABA dynamics. **(b)** Behavioral Paradigm. Mice underwent 30-minute sipper-access sessions while fiber photometry recordings were collected during consumption of water, a 50:50 Ensure-water mixture, or 0.1% saccharin solution presented on separate testing days. **(c)** Prior to testing, mice had ad libitum access to food and water in their home cages and were sequentially tested with water, Ensure, or saccharin. **(d)** Total lick counts increased with solution palatability, with Ensure eliciting the greatest number of licks across animals (n=14). Repeated-measures one-way ANOVA revealed a significant effect of solution (F(1.401, 18.21) = 56.74, p < 0.0001). Sidak’s multiple comparisons test: Ensure vs. Water. p < 0.0001; Ensure vs. Saccharin, p < 0.0001; Water vs. Saccharin, p= 0.0014. **(e)** Lick-aligned DLS GABA traces during consumption of water, Ensure, and saccharin. GABA fluorescence decreases following individual licking events, regardless of solution palatability. **(f)** Quantification of DLS GABA response as area under the curve (AUC). No significant effect of solution was detected (repeated-measures one-way ANOVA, F(2,12) = 0.9189, *p* = 0.4253; Dunnett’s multiple-comparisons test: Water vs. Ensure, *p* = 0.6202; Water vs. Saccharin, *p* = 0.3351). **(g)** Lick-aligned DMS GABA traces during consumption of water, Ensure, and saccharin. GABA fluorescence also decreases following licking, with response magnitude differing across solutions. **(h)** Quantification of DMS GABA responses as area under the curve (AUC). A significant effect of solution was observed (repeated-measures one-way ANOVA, F(2,12) = 4.320, *p* = 0.0386). Dunnett’s multiple-comparisons test revealed a significant reduction in GABA responses during Ensure consumption compared with water (*p* = 0.0228), whereas saccharin did not differ from water (*p* = 0.3258). The error bars represent S.E.M.s. ns=not significant, p>0.05, *p<0.05, **p<0.01, ****p<0.0001.

#### Food deprivation increases consumption of palatable liquids while delaying the return of DMS GABA signaling to baseline during consumption

Following the characterization of DMS GABA responses during consumption of palatable solutions with varying hedonic values in satiated animals, we next examined how DMS GABA signaling evolves over the course of continued intake within a session (see **Fig. 2a** for a schematic description). To quantify these within-session dynamics, we analyzed the GABA response across the initial 80 licks, grouped into bins of 10 licks. Comparison of the first ten licks (1–10) and the last ten licks (71–80) revealed that DMS GABA responses progressively increased over the course of consumption, eventually returning toward baseline during Ensure and saccharin intake, suggesting that GABA signaling is initially suppressed to permit consumption and gradually restored as intake continues (**Extended Data Fig. 2a,b**).

**Figure 2:**
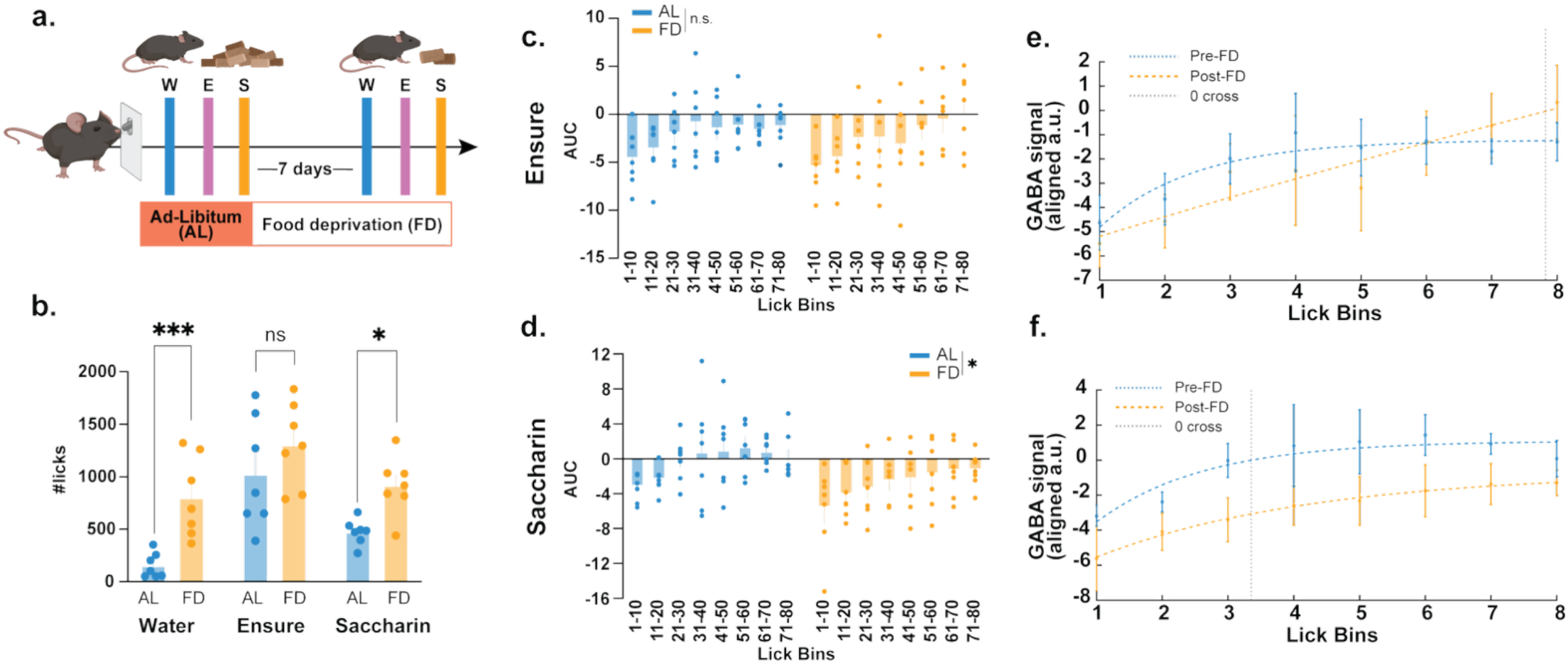
Food deprivation increases consumption of palatable liquids while delaying the return of DMS GABA signaling to baseline during consumption. **(a)** Experimental timeline. Mice (*n =* 7, 4 males and 3 females) were first tested during ad libitum (AL) feeding while consuming water, a 50:50 Ensure-water mixture, and 0.1% saccharin. Following 7 days of food deprivation (FD), the same mice were retested with each solution **(b)** Total lick counts during AL and FD conditions. Food deprivation significantly increased licking for water and saccharin, whereas Ensure consumption was not significantly altered. Repeated-measures two-way ANOVA revealed no solution x feeding interaction (F(2,12) = 2.137, *p* = 0.1607), but a significant main effect of feeding (F(1,6) = 29.61, *p* = 0.0016). Sidak’s multiple-comparisons test: Water, *p* = 0.0008; Ensure, *p* = 0.1403; Saccharin, *p* = 0.0132. **(c)** DMS GABA responses represented as area under the curve (AUC) across sequential lick bins during Ensure consumption. Food deprivation did not significantly alter lick-aligned GABA responses throughout the drinking bout (Repeated measures ANOVA, F(1,6) = 0.1609, *p* = 0.7022). **(d)** DMS GABA AUCs across sequential lick bins during saccharin consumption. Food deprivation significantly altered GABA accumulation during the drinking bout (Repeated measures ANOVA, F(1,6) = 6.240, *p* = 0.0467). **(e)** Nonlinear regression analysis of DMS GABA accumulation during Ensure consumption under ad libitum (AL) and food-deprived (FD) conditions. The fitted curves did not differ between feeding conditions (extra sum-of-squares F-test, F(3,106) = 0.883, p = 0.452), indicating that food deprivation did not significantly alter the dynamics of DMS GABA accumulation during Ensure consumption. **(f)** Nonlinear regression analysis of DMS GABA accumulation during saccharin consumption under ad libitum (AL) and food-deprived (FD) conditions. The fitted curves differed significantly between feeding conditions (extra sum-of-squares F-test, F(3,106) = 6.21, p = 0.0006), indicating that food deprivation altered the dynamics of DMS GABA accumulation during saccharin consumption. The error bars represent S.E.M.s. ns=not significant, p>0.05, *p<0.05, ***p<0.001.

To determine whether these dynamics are sensitive to motivational state, we retested the same animals after 7 days of food deprivation. Within-subject comparison revealed that food deprivation significantly increased saccharin consumption but not Ensure consumption (potentially due to a ceiling effect), relative to the same animals under ad libitum conditions (**Fig. 2b**). Notably, in both ad libitum and food-deprived conditions, DMS GABA responses showed a similar trajectory during Ensure consumption, characterized by an initial decrease during early licks followed by a gradual return toward baseline with continued intake (**Fig. 2c**), which is consistent with the comparable number of licks across feeding states for Ensure. In contrast, saccharin consumption revealed a divergence between feeding states. Under ad libitum conditions, DMS GABA responses increased from early (1–10) to late (71–80) licks, returning toward baseline within the session (**Fig. 2d; Extended Data Fig. 3a,b**). However, during food deprivation, this recovery was delayed and did not reach baseline within the same lick range (**Fig. 2d**), coinciding with increased saccharin consumption.

To directly compare the temporal evolution of GABA responses across conditions, we fit within-session trajectories with a nonlinear exponential model that captures the progressive recovery of GABA toward baseline across successive lick bins. This approach allowed us to quantify differences in the rate and extent of GABA recovery between feeding states. For Ensure consumption, the fitted trajectories did not differ between ad libitum and food-deprived conditions (**Fig. 2e**; Extra sum-of-squares F-test, p = 0.45), indicating that GABA recovery dynamics were preserved across motivational states. Consistent with this, the estimated time required for GABA signals to return toward baseline was comparable across conditions. In contrast, saccharin consumption crossed baseline (AUC=0) earlier than Ensure in ad-libitum conditions (∼71-80 lick bin for Ensure and 31-40 lick bin for saccharin) and revealed a significant difference in trajectory shape between feeding states (**Fig. 2f**; Extra sum-of-squares F-test, p = 0.0006), with food deprivation producing a rightward shift in the recovery curve, indicative of delayed GABA normalization. This delay corresponded to an extended period of suppressed GABA signaling during ongoing consumption. Together, these analyses confirm that while Ensure-evoked GABA dynamics remain stable across feeding states, saccharin responses exhibit a state-dependent delay in recovery, suggesting that prolonged suppression of DMS GABA is associated with increased intake under ad-libitum and food-deprived conditions.

#### DMS GABA suppression tracks and predicts lick bout initiation during consumption

To determine whether DMS GABA dynamics are temporally aligned with ongoing licking behavior, we examined the relationship between full session GABA fluctuations and lick bout occurrence. We first asked whether lick bouts preferentially occur during periods of reduced GABA signaling. Representative DMS GABA ΔF/F trace aligned to lick bouts, illustrating transient decreases in GABA that coincide with periods of active licking (**Fig. 3a**). Analysis of GABA values at lick onset revealed that lick bouts were significantly biased toward periods in which GABA levels were below baseline, indicating that bouts coincide with transient GABA dips (**Fig. 3b,c**).

**Figure 3.**
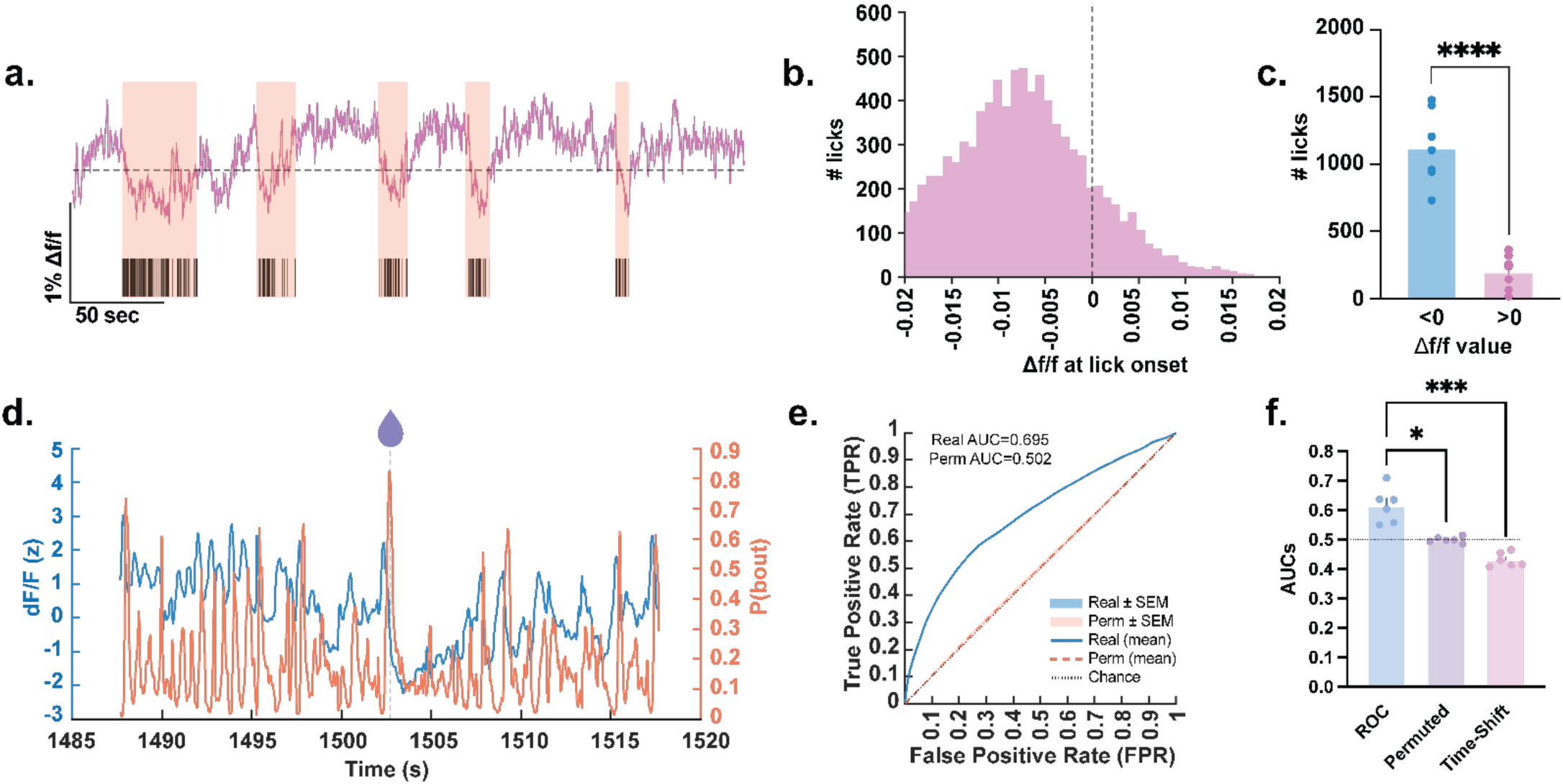
DMS GABA suppression tracks and predicts lick bout initiation during consumption. **(a)** Representative DMS GABA fiber photometry recording during consumption of an Ensure solution. Shaded regions indicate licking bouts. GABA fluorescence decreases during active licking and returns toward baseline as licking ceases. **(b)** Distribution of GABA ΔF/F values at lick onset, demonstrating that licking events predominantly occur during periods of reduced GABA fluorescence. **(c)** Quantification of lick frequency occurring when GABA ΔF/F was below or above zero (*n* = 7, 4 males and 3 females). Mice initiated significantly more licks when GABA ΔF/F was below zero than above zero (paired *t*-test, *t*(6) = 8.660, *p* = 0.0001). **(d)** Representative prediction from the leave-one-mouse-out logistic regression analysis. The model was trained on data from all remaining mice and evaluated on the completely held-out mouse. Model-generated lick-bout probability is shown together with the corresponding DMS GABA signal, illustrating the relationship between reduced GABA signaling and licking-related states around bout initiation. **(e)** Mean receiver operating characteristic (ROC) curves from leave-one-mouse-out cross-validation. For each fold, one mouse was held out completely for testing while the model was trained on all remaining mice. Mean ROC curves were calculated across the six held-out mice for the real model and control models (mean ROC AUC = 0.66; permuted ROC AUC = 0.50). **(f)** Quantification of ROC area under the curve (AUC) for the real, permuted-label, and time-shifted models across held-out mice (n = 6 mice). Each point represents the ROC AUC obtained from one held-out mouse. Repeated-measures ANOVA revealed a significant effect of model type (F(1.260, 6.300) = 44.42, *p* = 0.0003), with the trained model exhibiting significantly greater predictive performance than both control models (Dunnett post-hoc Real vs. Permuted p<0.05; Real vs. Time-shifted p<0.001). The error bars represent S.E.M.s. *p<0.05, ***p<0.001, ****p<0.0001.

To test whether DMS GABA fluctuations were predictive of licking-related states around bout initiation, we applied logistic regression with leave-one-mouse-out cross-validation. Models trained on all remaining animals reliably classified licking-related periods in the held-out mouse based on moment-to-moment DMS GABA dynamics (**Fig. 3d**). To determine whether this predictive relationship reflected temporally specific information in the GABA signal, performance was compared with permuted-label and time-shifted control models (**Fig. 3e,f**). The real model consistently outperformed both controls across held-out animals, demonstrating that DMS GABA dynamics contain temporally informative signals related to lick-bout initiation. Together, these findings show that transient decreases in DMS GABA are closely aligned with and predictive of consumption-related states, supporting a role for reduced DMS inhibitory signaling in sustaining ongoing Ensure consumption.

#### GLP-1 agonism reduces palatable reward consumption and enhances DMS GABA release

Next, we investigated whether pharmacological activation of GLP-1 receptors alters DMS GABA modulation during consumption of Ensure. To diminish the reward value of Ensure, we administered the GLP-1 receptor agonist semaglutide at 0.026 mg/kg^34,36,37^ subcutaneously 1 hour prior to Ensure access for 3 consecutive days. After a 14-day washout period, the same 3-day procedure was repeated with 0.9% saline (vehicle) injections (see **Fig. 4a** for a schematic description). Semaglutide decreased lick counts compared to saline days (**Fig. 4b**; **Extended Data Fig. 4a**) and induced significant body weight loss in the animals (**Fig. 4c**). Furthermore, semaglutide administration significantly reduced the number of licks (**Fig. 4d**) and lick bouts (**Fig. 4e**), produced a trend toward shorter lick bout duration (**Fig. 4f**), and reduced total consumption time (**Fig. 4g**) compared to the first day of saline injections. Finally, locomotor analysis during reward consumption sessions revealed that semaglutide reduced total distance traveled in the chamber, while movement velocity remained unchanged (**Extended Data Fig. 4b,c**), suggesting that this effect is unlikely to reflect a motor deficit and is instead attributable to reduced motivation to engage in reward-seeking behavior.

**Figure 4.**
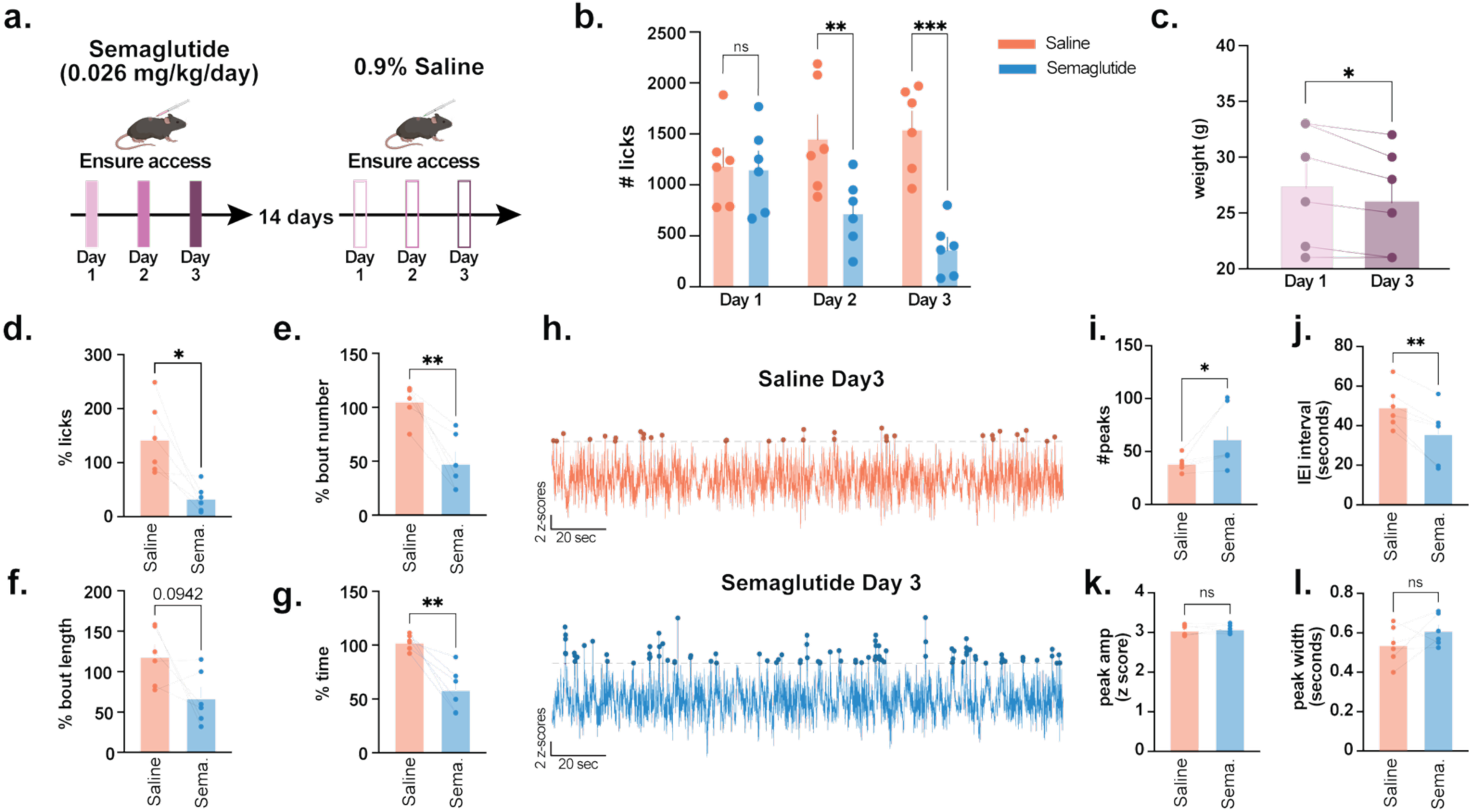
Repeated semaglutide administration reduces palatable liquid intake while increasing DMS GABA event frequency during consumption. **(a)** Experimental design. Mice (*n* = 6, 3 males and 3 females) completed three consecutive days of 30-min Ensure access sessions following subcutaneous injections of semaglutide (0.026 mg/kg/day). After a 14-day washout period, the same mice underwent an identical testing protocol following 0.9% saline injections. **(b)** Total lick counts across the three testing days during saline and semaglutide treatment showing decreased Ensure consumption after 3 days of semaglutide injections but not after saline injections (Repeated-measures two-way ANOVA revealed a significant drug × day interaction F(2,10) = 10.81, *p* = 0.0032 and a significant main effect of drug F(1,5) = 8.457, *p* = 0.0335. Sidak’s multiple-comparisons test: Day 1, *p* > 0.05; Day 2, *p* = 0.0056; Day 3, *p* = 0.0002). **(c)** Body weight before (Day 1) and after (Day 3) semaglutide treatment. Semaglutide significantly reduced body weight following three days of treatment (paired *t*-test, *t*(5) = 3.162, *p* = 0.0250). **(d)** Total lick counts on Day 3 expressed as a percentage of saline Day 1 values. Semaglutide significantly reduced overall consumption (paired *t*-test, *t*(5) = 3.428, *p* = 0.0187). **(e)** Number of licking bouts during Day 3 expressed as a percentage of saline Day 1 values. Semaglutide significantly reduced the total number of licking bouts (paired *t*-test, *t*(5) = 5.353, *p* = 0.0031). **(f)** Lick bout length on Day 3 expressed as a percentage of saline Day 1 values. Although semaglutide decreased the bout length duration, the effect did not reach statistical significance (paired *t*-test, *t*(5) = 2.062, *p* = 0.0942). **(g)** Time from the first to the last lick during Day 3 expressed as a percentage of saline Day 1 values. Semaglutide significantly decreased the duration of the drinking session (paired *t*-test, *t*(5) = 5.072, *p* = 0.0039). **(h)** Representative DMS GABA fluorescence traces illustrating detected GABA events during Ensure consumption under saline (top) and semaglutide (bottom) conditions. Peak detection was used to quantify event frequency, inter-event interval, amplitude, and width. **(i)** GABA event frequency during Ensure consumption. Semaglutide significantly increased DMS GABA event frequency compared with saline (Wilcoxon matched-pairs signed-rank test, *W* = 21, *p* = 0.0312). **(j)** Mean inter-event interval (IEI) between DMS GABA events. Semaglutide significantly reduced the interval between events (paired *t*-test, *t*(5) = 5.226, *p* = 0.0034). **(k)** Mean DMS GABA event amplitude. Semaglutide did not significantly alter event amplitude (paired *t*-test, *t*(5) = 0.6411, *p* = 0.5497). **(l)** Mean DMS GABA event width. Semaglutide did not significantly alter event width (paired *t*-test, *t*(5) = 1.494, *p* = 0.1953). The error bars represent S.E.M.s. ns=not significant, p>0.05, *p<0.05, **p<0.01.

Next, we examined how repeated semaglutide administration affects DMS GABA release dynamics by comparing saline Day 3 and semaglutide Day 3 sessions within the same animals. Notably, semaglutide altered baseline DMS GABA dynamics reflected by an increased number of peaks and decreased inter-peak intervals, without affecting peak amplitude or width (**Fig. 4h–l**), suggesting a shift toward more sustained inhibitory signaling.

Examining lick-coupled GABA release, semaglutide did not alter GABA transients early in the session (first 5 licks), but these effects emerged later in the session (**Fig. 5a,b**). During late-session consumption (licks 71–80), semaglutide increased DMS GABA signals time-locked to licking (**Fig. 5c,d**). To account for variability in total lick counts across animals and align GABA dynamics to relative session progression, we also analyzed signals across lick quartiles. Both semaglutide and saline exhibited progressively increasing DMS GABA transients across quartiles (**Fig. 5e–i**); however, this increase was significantly greater during the fourth quartile following semaglutide (**Fig. 5i**). Semaglutide also accelerated the recovery of DMS GABA toward baseline across lick bins, producing a steeper trajectory relative to saline controls (**Fig. 5j**). Consistent with this, semaglutide shifted the GABA trajectory leftward (**Fig. 5k**), indicating an earlier emergence of elevated GABA signaling during ongoing consumption. Together, these findings suggest that semaglutide enhances the progressive recruitment of DMS GABA signaling during ongoing consumption, consistent with a strengthening inhibitory “brake” on behavior.

**Figure 5:**
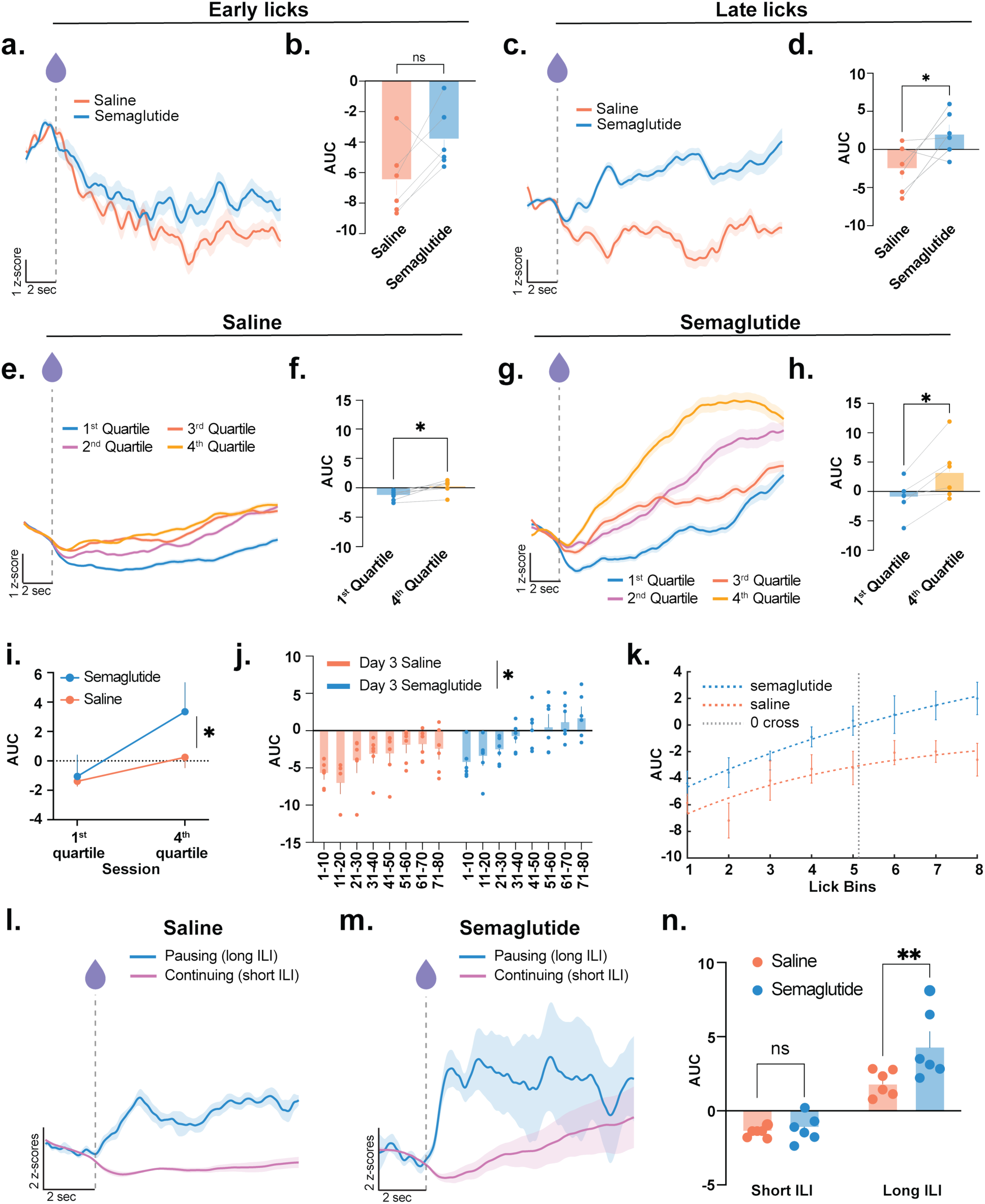
Repeated semaglutide administration enhances DMS GABAergic rebound during late phases of palatable liquid consumption. **(a)** Average lick-aligned DMS GABA fluorescence traces aligned to early licking events during Ensure consumption following saline (red) or semaglutide (blue) treatment. Early licking elicited comparable GABA responses between treatment groups. **(b)** Quantification of early-lick GABA responses (AUC). Semaglutide did not significantly alter early-lick GABA activity compared with saline (paired *t*-test, *t*(5) = 2.296, *p* = 0.0701; *n* = 6, 3 males and 3 females). **(c)** Average peri-lick DMS GABA fluorescence traces aligned to late licking events during Ensure consumption following saline and semaglutide treatment. Semaglutide-treated mice exhibited a more pronounced post-lick increase in DMS GABA fluorescence during late consumption. **(d)** Quantification of late-lick GABA responses (AUC). Semaglutide significantly increased late-lick GABA activity relative to saline (paired *t*-test, *t*(5) = 2.617, *p* = 0.0473). **(e)** Average peri-lick DMS GABA fluorescence traces separated into four sequential lick quartiles during Day 3 saline treatment. Traces illustrate the progression of GABA responses as the drinking bout advances from the first to the fourth quartile. **(f)** Comparison of first and fourth lick quartiles during saline treatment. GABA responses were significantly greater during the fourth quartile than the first (paired *t*-test, *t*(5) = 3.950, *p* = 0.0109). **(g)** Average peri-lick DMS GABA fluorescence grouped by lick quartile during Day 3 semaglutide treatment, illustrating progressive enhancement of GABA signaling as consumption continued. **(h)** Comparison of first and fourth lick quartiles during semaglutide treatment. GABA responses increased significantly between the first and fourth quartiles (paired *t*-test, *t*(5) = 3.441, *p* = 0.0184). **(i)** Comparison of first-and fourth-quartile GABA responses between saline and semaglutide treatment. Repeated-measures two-way ANOVA revealed no significant interaction (F(1,5) = 4.342, *p* = 0.0916), but significant main effects of drug (F(1,5) = 19.74, *p* = 0.0067). Sidak’s multiple-comparisons test identified a significant increase during the fourth quartile (*p* = 0.0426), whereas first-quartile responses did not differ (*p* > 0.05). **(j)** Mean DMS GABA responses across sequential lick bins during Day 3. Repeated-measures two-way ANOVA revealed no drug × lick-bin interaction (F(3.013,15.07) = 0.7558, *p* = 0.5365), but significant main effects of drug (F(1,5) = 9.123, *p* = 0.0294) and lick number (F(1.596,7.978) = 19.97, *p* = 0.0011). **(k)** Nonlinear exponential regression of DMS GABA accumulation during Ensure consumption. Regression curves differed significantly between saline and semaglutide conditions (extra sum-of-squares F-test, F(3,90) = 10.85, *p* < 0.0001). **(l)** Average DMS GABA fluorescence aligned to licking events preceding short inter-lick intervals (ILIs; continuous licking) and long ILIs (lick pauses ≥5 s) during saline treatment. **(m)** Average DMS GABA fluorescence aligned to licking events preceding short and long inter-lick intervals during semaglutide treatment. Long ILIs were preceded by larger increases in DMS GABA fluorescence than short ILIs. **(n)** Quantification of GABA responses preceding short and long ILIs. Repeated-measures two-way ANOVA revealed a significant interaction between drug and ILI duration (F(1,5) = 9.261, *p* = 0.0286), with significant main effects of drug (F(1,5) = 6.655, *p* = 0.0495) and interval length (F(1,5) = 66.30, *p* = 0.0005). Sidak’s multiple-comparisons test showed no difference for short ILIs (*p* > 0.05), whereas long ILIs exhibited significantly greater GABA responses following semaglutide treatment (*p* = 0.0097). The error bars represent S.E.M.s. ns=not significant, p>0.05, *p<0.05, **p<0.01.

To further test the hypothesis that DMS GABA functions as an inhibitory “brake” on consumption, we examined whether GABA signaling increases prior to pauses in licking behavior. If GABA constrains ongoing intake, we expect larger GABA responses to precede longer inter-lick intervals (ILIs), reflecting transitions out of active consumption. We defined short ILIs as <0.5 s (continuous licking) and long ILIs as >5 s (pausing). By day 3, semaglutide selectively enhanced DMS GABA signals preceding long ILIs, but not short ILIs (**Fig. 5l–n**), indicating an increased inhibitory signal prior to consumption pauses. Consistent with this, semaglutide disrupted the coupling between GABA decreases and lick initiation (**Fig. 6a**) and reduced the likelihood of licks occurring during periods of below-baseline GABA signaling (**Fig. 6b,c**). Thus, semaglutide strengthens GABA-mediated inhibitory control over consumption and abolishes the permissive role of GABA dips in predicting lick initiation.

**Figure 6.**
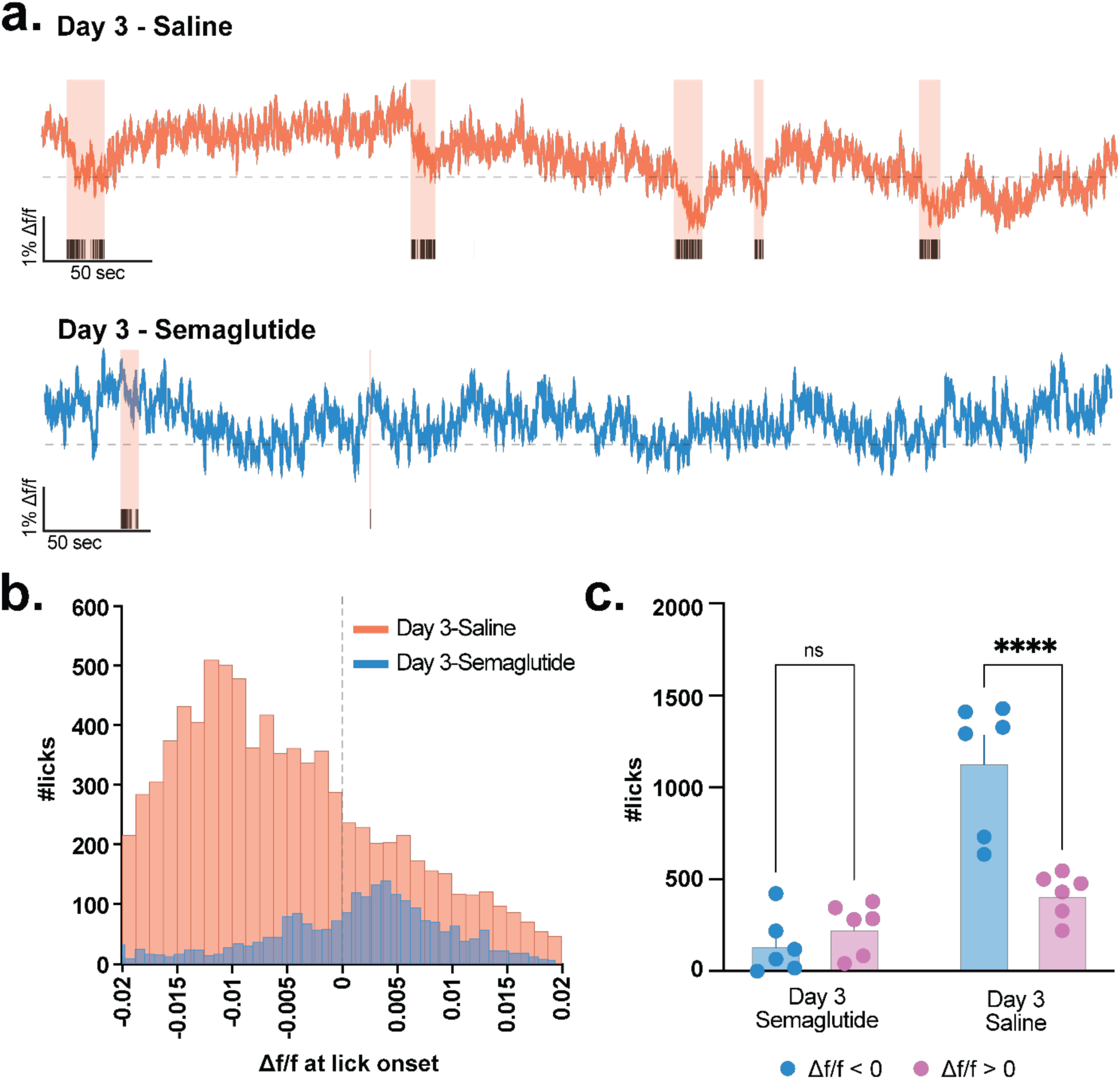
Semaglutide disrupts the temporal coupling between DMS GABA dynamics and licking behavior. **(a)** Representative DMS GABA fluorescence recordings during Ensure consumption following Day 3 saline (top) or semaglutide (bottom) treatment. Shaded regions indicated licking bouts. Under saline treatment, licking bouts predominantly occurred during decreases in DMS GABA fluorescence, whereas this temporal relationship was disrupted following semaglutide treatment. **(b)** Distribution of DMS GABA ΔF/F values at lick onset during Day 3 saline and semaglutide treatment. Lick onsets during saline were enriched during periods of negative GABA fluorescence, whereas semaglutide shifted the distribution toward baseline. **(c)** Quantification of lick counts occurring when DMS GABA fluorescence was below (ΔF/F < 0) or above baseline (ΔF/F > 0) during Day 3 saline and semaglutide treatment (*n* = 6, 3 males and 3 females). Lick bouts mostly occurred when the GABA signal was below 0 after saline injections, but this effect was abolished after repeated semaglutide injections (Repeated-measures two-way ANOVA: treatment x GABA state interaction F(1,5) = 6.144, *p* = 0.0559, and the main effect of treatment F(1,5) = 26.58, *p* = 0.0036). Sidak’s multiple-comparisons test showed no difference between GABA states following semaglutide treatment (*p* > 0.05), whereas saline-treated mice exhibited significantly more licks during periods of negative GABA fluorescence than positive GABA fluorescence (*p* < 0.0001).The error bars represent S.E.M.s. ns=not significant, p>0.05, ****p<0.0001.

Together, these findings demonstrate that GLP-1 receptor activation alters DMS GABA dynamics during consumption of Ensure, promoting sustained inhibitory signaling that suppresses consumption and disrupts the permissive role of GABA dips in driving reward-seeking behavior.

#### Repeated semaglutide enhances coordinated DMS neuronal ensemble activity at baseline conditions

Our photometry results suggested that repeated semaglutide administration enhances DMS GABA signaling at baseline. We therefore hypothesized that glucagon-like peptide-1 receptor (GLP-1R) agonism would alter the organization of DMS neuronal ensembles. To test this, we recorded calcium activity from the same DMS neuronal populations before and after repeated semaglutide treatment. Mice received AAV-GCaMP (AAV1.CaMK2a.GCaMP6m.WPRE.SV40) infusions into the DMS and were implanted with GRIN lenses 6 weeks before imaging experiments (see **Extended Data Fig. 1b** for GRIN lens placements). Single-cell calcium activity was recorded under baseline (freely moving) conditions following three consecutive days of systemic saline or semaglutide (0.026 mg/kg, i.p.) administration (**Fig. 7a,b,c**). Consistent with previous reports, semaglutide-treated mice exhibited significant weight loss (∼5%) after the third injection (**Extended Data Fig. 5**).

**Figure 7.**
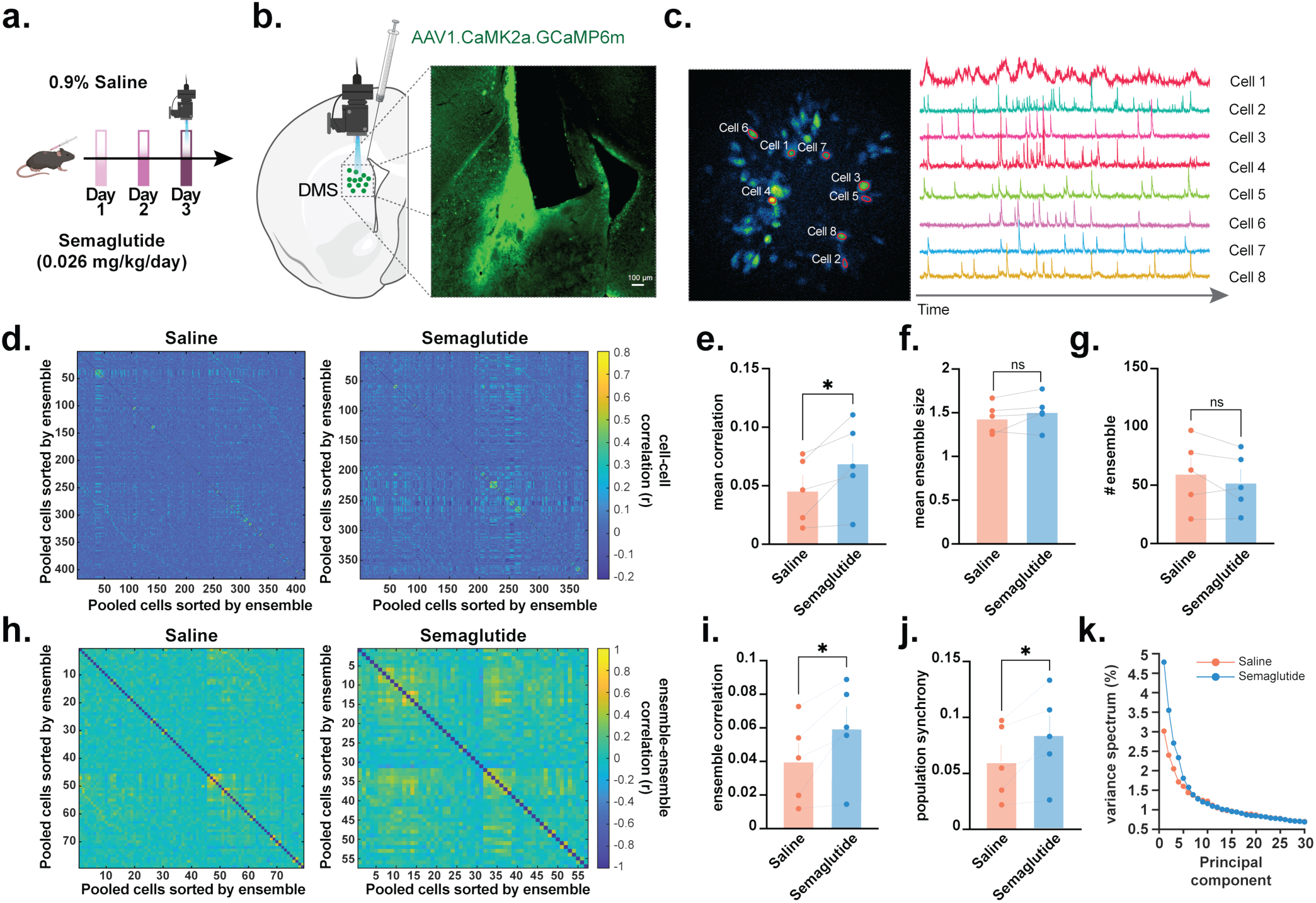
Repeated semaglutide enhances coordinated DMS neuronal ensemble activity at baseline conditions. **(a)** Experimental schematic. Mice (*n* = 5, 3 males and 2 females) underwent three consecutive days of saline or semaglutide treatment before calcium imaging. **(b)** AAV1.CaMK2a.GCaMP6m.WPRE.SV40 was expressed in DMS neurons. Representative histology image showing GCaMP expression and GRIN lens placement in the DMS. **(c)** Representative miniscope field of view (left) showing identified DMS neurons used for analysis and corresponding calcium fluorescence traces from individual neurons (right). **(d)** Pairwise ensemble-organized matrices during saline (left) and semaglutide (right) treatment. Cells are ordered by ensemble membership, illustrating patterns of correlated activity across the recorded neuronal population. **(e)** Mean pairwise correlation coefficient across all recorded neurons. Semaglutide significantly increased coordinated neuronal activity (paired *t*-test, *t*(4) = 2.909, *p* = 0.0437). **(f)** Semaglutide did not significantly alter the number of neurons participating within individual ensembles (paired *t*-test, *t*(4) = 1.484, *p* = 0.2120). **(g)** Semaglutide did not significantly change the number of ensembles identified in freely moving mice (paired *t*-test, *t*(4) = 1.382, *p* = 0.2393). **(h)** Pairwise ensemble-organized matrices during saline (left) and semaglutide (right) treatment. Neurons are ordered by hierarchical clustering such that neurons assigned to the same functional ensemble are grouped together, revealing within-and between-ensemble correlation structure. **(i)** Semaglutide significantly increased correlations among neurons belonging to different ensembles compared with saline treatment (paired t-test, t(4) = 3.084, p = 0.0368). **(j)** Population synchrony across the recorded neuronal population. Semaglutide significantly increased overall network synchrony (paired *t*-test, *t*(4) = 2.875, *p* = 0.0452). **(k)** Principal component variance spectrum of DMS population activity under saline and semaglutide treatment. Semaglutide shifted the variance explained toward the first principal components, indicating that a larger proportion of population activity could be captured by fewer dimensions. The error bars represent S.E.M.s. ns=not significant, p>0.05, *p<0.05.

Analysis of individual neuronal activity revealed that semaglutide did not alter either the frequency or amplitude of calcium transients (**Extended Data Fig. 6**), suggesting that overall neuronal excitability remained largely unchanged. In contrast, semaglutide produced marked changes in network organization. Pairwise cell-cell correlations were significantly increased following semaglutide treatment (**Fig. 7d,e**). Ensemble analysis further demonstrated that, although the number and size of neuronal ensembles were unchanged (**Fig. 7,f,g**), ensemble-to-ensemble correlations and overall population synchrony were significantly enhanced after semaglutide administration (**Fig. 7j**). Principal component analysis additionally revealed that neural population activity occupied a lower-dimensional space after semaglutide treatment, with a smaller number of principal components accounting for a greater proportion of the total variance (**Fig. 7k, Extended Data Fig. 7**), indicating a more coordinated population state.

Together, these findings demonstrate that repeated GLP-1R agonism reorganizes DMS network activity without substantially altering the activity of individual neurons. Combined with our photometry results showing enhanced DMS GABA release, these data suggest that semaglutide promotes a more synchronized and constrained DMS network state, consistent with increased GABAergic regulation of DMS output.

#### Optogenetic stimulation of GABAergic interneurons partially mimics the effects of GLP-1R agonism on reward consumption

Next, we asked whether GABA release from DMS GABAergic interneurons is sufficient to recapitulate the behavioral effects of semaglutide by optogenetically stimulating local GABAergic interneurons (GINs). We injected AAV.PHP.eB-hDLX-minBG-iCre-4X2C-WPRE3-BGHpA to drive Cre-recombinase expression broadly across GINs, enabling pan-GIN targeting^38^, as well as either a Cre-dependent excitatory opsin (AAV5-Syn-FLEX-rc[ChrimsonR-tdTomato]; *Chrimson* animals) for optogenetically stimulated excitation or a control virus (AAV5-hSyn-DIO-mCherry; *mCherry* animals) (**Fig. 8a**; see **Extended Data Fig. 1c** for optogenetic fiber implant placements). Chrimson expression in Cre-expressing GINs was verified histologically in the DMS (**Extended Data Fig. 8** or **Extended Data Fig. 9a**). All optogenetic animals were also injected with iGABA.Sn.FR2 (pGP-AAV5-syn-iGABA.Sn.FR2; **Fig. 9a**) to validate DMS GABA release during laser stimulation (10Hz, 5mW) from GINs (**Fig. 8b,c**).

**Figure 8.**
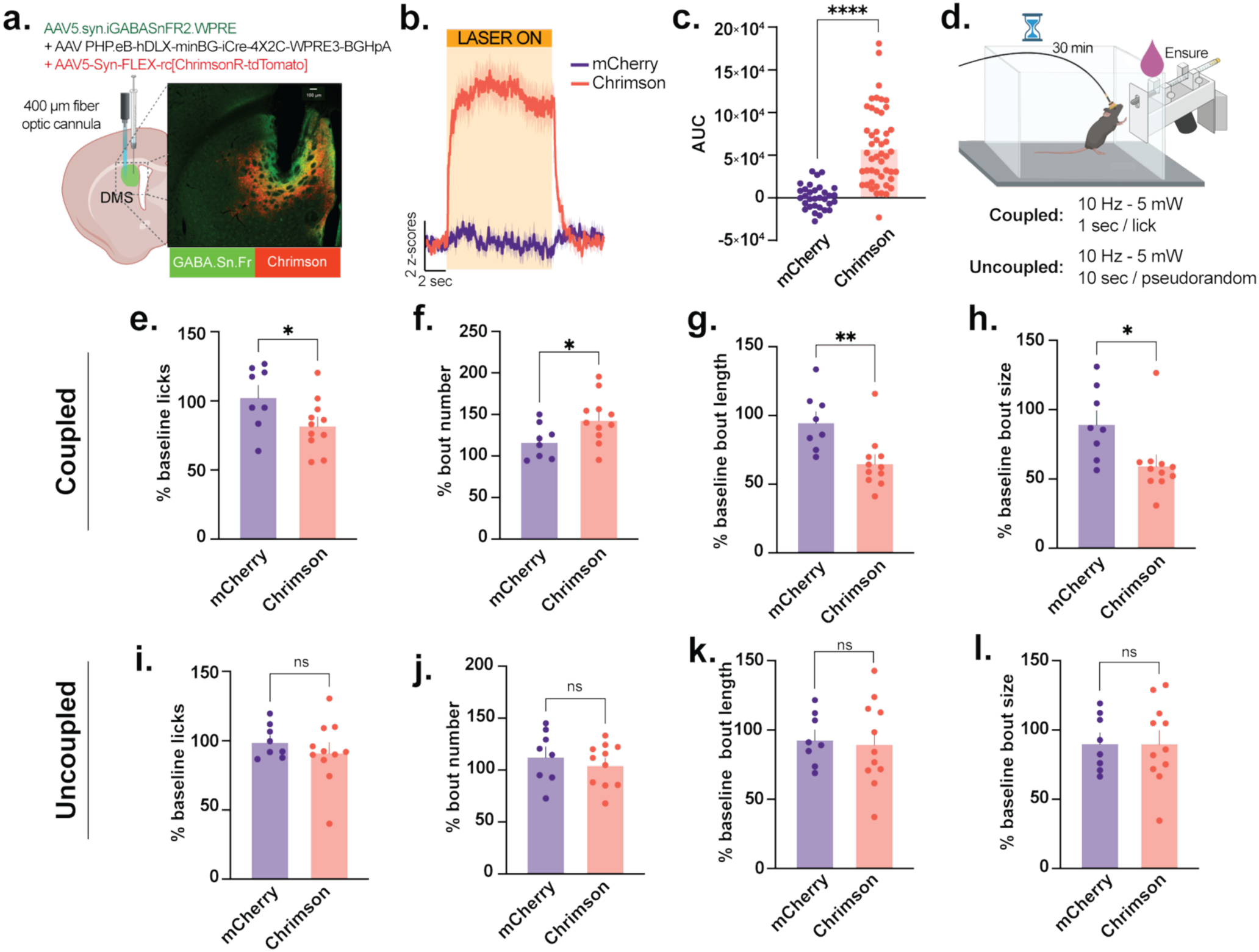
Optogenetic activation of DMS GABAergic interneurons suppresses reward consumption. **(a)** Experimental strategy. Mice received unilateral injections of AAV5-syn-iGABA.Sn.FR2 together with either AAVPHP.eB-hDLX-minBG-iCre-4X2C-WPRE3-BGHpA and AAV5-Syn-FLEX-ChrimsonR-tdTomato (Chrimson; *n* = 11, 8 males and 3 females) or AAV5-hSyn-DIO-mCherry (mCherry; *n* = 8, 4 males and 4 females) into the DMS, followed by implantation of a 400-µm fiber-optic cannula. This approach enabled simultaneous fiber photometry recordings of extracellular GABA and optogenetic activation of DMS GABAergic interneurons. Representative histology confirms viral expression of both viruses. **(b)** Representative peri-stimulation GABA fluorescence traces following 589-nm laser stimulation (5 mW, 10 Hz) in Chrimson (red) and mCherry (blue) mice. Optogenetic stimulation evoked robust increases in extracellular GABA selectively in Chrimson-expressing mice. **(c)** Quantification of stimulation-evoked GABA responses (AUC). Chrimson mice exhibited significantly greater GABA release following laser stimulation than mCherry controls (unpaired *t*-test, *t*(78) = 6.918, *p* < 0.0001, n=32-48 stimulations). **(d)** Behavioral paradigm. Following photometry validation, mice underwent 30-minute Ensure access sessions. Once reliable, Ensure consumption was established; mice were given a session in which each lick was paired with laser stimulation (589 nm; 5 mW, 10 Hz, 1 s), as well as the Ensure delivery. After three recovery sessions without stimulation, mice underwent an uncoupled condition in which laser stimulation (589 nm; 5 mW, 10 Hz, 10 s) was delivered pseudorandomly and independently of licking behavior. **(e)** Percentage of baseline licks during lick-coupled optogenetic stimulation. Chrimson mice exhibited a greater reduction in licking than mCherry controls (unpaired *t*-test, *t*(17) = 2.165, *p* = 0.0449). **(f)** Percentage of baseline licking bouts during coupled stimulation. Chrimson mice displayed significantly more licking bouts than mCherry controls (unpaired *t*-test, *t*(17) = 2.218, *p* = 0.0405). **(g)** Percentage of baseline bout length during coupled stimulation. Bout length was significantly reduced in Chrimson mice compared with mCherry controls (unpaired *t*-test, *t*(17) = 3.166, *p* = 0.0056). **(h)** Percentage of baseline bout size during coupled stimulation. Chrimson mice exhibited significantly fewer licks per bout than mCherry controls (unpaired *t*-test, *t*(17) = 2.603, *p* = 0.0186). **(i)** Percentage of baseline licks during pseudorandom (uncoupled) optogenetic stimulation. No significant difference was observed between Chrimson and mCherry mice (unpaired *t*-test, *t*(17) = 0.8652, *p* = 0.3990) **(j)** Percentage of baseline licking bouts during uncoupled stimulation. The number of licking bouts did not differ between groups (unpaired *t*-test, *t*(17) = 0.7888, *p* = 0.4411). **(k)** Percentage of baseline bout length during uncoupled stimulation. Bout length was not significantly altered by uncoupled stimulation (unpaired *t*-test, *t*(17) = 0.2473, *p* = 0.8076). **(l)** Percentage of baseline bout size during uncoupled stimulation. Bout size did not differ between Chrimson and mCherry mice (unpaired *t*-test, *t*(17) = 0.0094, *p* = 0.9926). The error bars represent S.E.M.s. ns=not significant, p>0.05, *p<0.05, **p<0.01, ****p<0.0001.

Activation of DMS GABAergic interneurons did not induce detectable locomotor or aversive side effects. In open-field testing (**Extended Data Fig. 9a**), Chrimson animals did not demonstrate any difference in immobility, velocity, or distance traveled (**Extended Data Fig. 9b,c,d**) compared to *mCherry* animals, suggesting that stimulation of DMS GINs does not alter locomotion. Similarly, the conditioned place aversion (CPA) test (**Extended Data Fig. 10a**) did not reveal any difference in chamber preference (**Extended Data Fig. 10b,c**), indicating that stimulation of DMS GINs does not intrinsically induce aversion.

Finally, we tested whether optogenetic activation of DMS GABAergic interneurons, thereby elevating DMS GABA release, alters reward consumption. When laser stimulation (1 s) was coupled to each lick, *Chrimson* animals exhibited reduced total lick counts (**Fig. 8e**) and increased lick bout frequency (**Fig. 8f**), accompanied by shorter and smaller lick bouts compared to *mCherry* controls (**Fig. 8g,h**), indicating that GIN-driven GABA release promotes disengagement from ongoing consumption. To determine whether these effects depend on temporal coupling to behavior, we delivered laser stimulation in a pseudo-random manner. Under these conditions, uncoupled GABA elevation did not alter consumption measures between *Chrimson* and *mCherry* animals (**Fig. 8i–l**), demonstrating that the effects of DMS GABA release on reward consumption are time-locked to ongoing behavior.

Together, these findings demonstrate that optogenetic activation of DMS GABAergic interneurons is sufficient to disengage animals from ongoing consumption by shortening lick bouts and reducing intake but does not fully recapitulate the motivational suppression induced by semaglutide.

## Discussion

In the present study, we demonstrate that dorsomedial striatal GABA signaling dynamically tracks the motivational and hedonic value of palatable rewards throughout consumption. Unlike the dorsolateral striatum, DMS GABA responses scaled with reward palatability and showed a rapid decrease at lick onset, followed by a gradual recovery as consumption continued. Conditions that increased motivation to consume, such as food deprivation, delayed this recovery, whereas the semaglutide-induced decrease in consumption accelerated GABA recovery, increased inhibitory signaling during consumption, and weakened the relationship between GABA suppression and individual licking events. Semaglutide also shifted DMS network activity toward a more synchronized and less variable state, suggesting altered population-level encoding of reward. Finally, optogenetic activation of DMS GABAergic interneurons reduced lick bout duration and overall consumption, demonstrating that increased DMS inhibitory signaling is sufficient to suppress consummatory behavior. Together, these findings identify DMS GABA signaling as a dynamic inhibitory mechanism that links reward value to consummatory behavior and contributes to transitions between consumption engagement and disengagement.

Prior work has established that the dorsal striatum contributes to food-directed action and consumption, with the DMS supporting flexible reward-guided behavior and food intake, and the DLS becoming increasingly engaged during habitual or binge-like consumption of palatable foods.^39,40^ Other studies have shown that dorsal striatal activity changes during food approach and consumption, and that neuromodulatory signals, such as enkephalin in the anteromedial dorsal striatum, can promote intense intake of palatable food.^41,42^ Building on this literature, our studies identify DMS GABAergic signaling as a previously unresolved, subregion-specific neurochemical mechanism linking palatability to consummatory behavior. We show that GABA release in the DMS exhibits rapid, consumption-coupled decreases during intake of water, saccharin, and Ensure, a pattern that is largely absent in the DLS, indicating subregion-specific encoding. Notably, these transient GABA decreases are temporally aligned with licking behavior and predict licking-related states around bout initiation, suggesting that DMS GABA dynamics provide a temporally structured signal associated with ongoing consumption. Thus, this work extends prior dorsal striatal feeding studies by moving beyond lesion, neuronal activity, and opioid-based mechanisms to reveal real-time GABA release in the DMS as a novel signal for palatability-driven consumption.

Our results further show that motivational state modulates DMS GABAergic encoding of palatable consumption. This fits with prior work implicating the DMS in goal-directed reward seeking and incentive-value-guided action, including studies showing that DMS function is required for instrumental action-outcome learning and that DMS activity tracks expected return and performance vigor^43,44^. Food deprivation increased water and saccharin consumption. This altered licking dynamics across bouts, consistent with enhanced incentive value^45,46^. These behavioral changes were paralleled by shifts in DMS GABA signaling, supporting the idea that internal state gates the striatal encoding of palatability. Notably, because dorsal striatal opioid/enkephalin signaling has been shown to promote intense consumption of palatable foods, our findings extend this literature by identifying DMS GABA release as a temporally precise signal that tracks ongoing intake^41^. The deprivation-related shift in DMS GABA dynamics may reflect a delayed inhibitory “brake” or prolonged engagement window, allowing palatable intake to remain motivationally salient before satiety-related feedback suppresses further consumption. Together, these findings position the DMS as a flexible node that integrates hedonic value, incentive salience, and satiety state to guide goal-directed consumption^47,48^.

We further demonstrate that GLP-1 receptor agonism suppresses intake and fundamentally alters DMS GABA dynamics. Consistent with these changes in GABA signaling, repeated semaglutide administration also reorganized DMS neural ensemble activity without altering the frequency or amplitude of individual neuronal calcium events. Instead, semaglutide increased cell-cell correlations, strengthened ensemble synchrony, and reduced the dimensionality of population activity, suggesting a shift toward a more coordinated network state. These findings build on previous work demonstrating that central GLP-1 receptor signaling suppresses food reward and motivated intake through mesolimbic circuits, including the VTA and nucleus accumbens^34,36,49^. Previous work has established that GLP-1 signaling regulates both homeostatic and hedonic aspects of feeding, with GLP-1 receptor activation decreasing palatable food intake and food-directed motivation through distributed neural circuits, including the lateral parabrachial nucleus and VTA^32,49,50^. More recent studies have further identified discrete central circuits through which GLP-1R agonists suppress hedonic aspects of feeding, highlighting the distributed neural networks through which GLP-1 regulates food reward^51^. Our findings extend this emerging framework by identifying DMS GABA signaling as a potential downstream mechanism through which GLP-1 agonism reshapes the temporal dynamics of ongoing consumption.

Consistent with this interpretation, semaglutide reduced licking and disrupted the temporal coupling between GABA decreases and lick onset, indicating a breakdown in consumption-related encoding. This finding is consistent with evidence that GLP-1R activation can recruit inhibitory GABAergic circuitry to suppress reward seeking. A recent study showed that GLP-1R signaling in midbrain circuits reduces cocaine seeking through engagement of GABAergic projection to VTA.^52,53^ However, our results are novel in showing that GLP-1 agonism alters GABA release dynamics within the dorsal striatum during natural reward consumption, rather than only modulating mesolimbic drug-seeking circuits. Because GLP-1 receptors are expressed in striatal regions and have been reported in GABAergic neuronal populations, semaglutide may act directly or indirectly on local DMS circuits, potentially through GLP-1R-expressing GABAergic interneurons or differential modulation of D1-and D2-expressing MSNs^54,55^. The observation that network organization changed despite stable single-neuron activity further suggests that GLP-1 signaling primarily influences the coordination of DMS circuits rather than the excitability of individual neurons. At the same time, GLP-1 agonist enhanced GABAergic rebound signals preceding pauses in licking, particularly during later stages of consumption and before longer inter-lick intervals. These results suggest that GLP-1 does not simply reduce reward value, but instead reshapes the temporal structure of striatal activity, weakening signals that sustain intake while strengthening those associated with satiety-like behavioral termination.

Across conditions, our data support a model in which DMS GABA signaling reflects a dynamic between engagement and disengagement states during consumption. Decreases in GABA are associated with ongoing licking, whereas increases precede pauses and cessation, suggesting that local inhibitory signaling may help structure the timing of when animals initiate, sustain, or terminate consummatory bouts. This interpretation fits with prior work showing that GLP-1 receptor activation suppresses food intake and food reward through mesolimbic and accumbal circuits^34,36,49^. Importantly, GLP-1 has also been linked to GABAergic control of motivated behavior. GLP-1 receptor activation in the VTA and related brainstem-midbrain circuits reduces cocaine seeking by recruiting GABAergic inhibition of dopamine neurons^37,52,53^. Our results extend this GLP-1-GABA framework to natural reward consumption by showing that semaglutide shifts DMS GABA dynamics toward disengagement, weakening lick-coupled signals that sustain intake while strengthening GABA rebounds that precede pauses. Because dorsal striatal circuits are central for translating motivation into action, this GLP-1R agonist-induced GABA shift may represent a circuit mechanism through which reduced reward value is converted into altered motor output, shorter licking bouts, and earlier satiety-like behavioral termination. Consistent with this model, optogenetic manipulation altered licking behavior and bout structure, supporting a causal role for dorsal striatal GABAergic interneuron-driven signaling in regulating consummatory actions.

Finally, our single-cell calcium imaging data suggest that enhanced GABA signaling is accompanied by increased coordination and reduced dimensionality of DMS neural ensembles following repeated semaglutide administration. Because DMS population activity is thought to support flexible action selection and action-outcome representations^6,56,57^, this more constrained network state may limit the dynamic neural computations necessary to sustain ongoing reward consumption. Consistent with this idea, optogenetic stimulation of GABAergic interneurons decreased consumption and altered bout structure, partially mimicking semaglutide’s suppressive effects. However, this manipulation appeared to primarily affect the motor organization of licking, suggesting that local DMS GABAergic signaling may be sufficient to interrupt or terminate ongoing intake but not fully reproduce the broader motivational decrease produced by GLP-1 receptor agonism. Because GLP-1 signaling is known to reduce food reward through mesolimbic and accumbal circuits, the “missing” motivational component may arise upstream or in parallel through VTA-nucleus accumbens, hypothalamic, and hindbrain circuits that regulate incentive salience, satiety, and reward valuation^34,36,49^. Together, these findings support a novel model in which motivational state and GLP-1 signaling shape reward consumption through distributed reward and metabolic circuits, while DMS GABA dynamics provide a temporally precise output mechanism that gates the transition from consumption engagement to disengagement.

Future studies using cell-type-and projection-specific approaches will be critical for defining how distinct striatal circuits contribute to the balance between consumption and cessation. This is especially important because GLP-1 receptor agonists are now clinically established for obesity treatment, with semaglutide producing substantial body-weight reduction in adults with overweight or obesity^58^. Beyond obesity, growing preclinical and clinical evidence suggests that GLP-1 signaling may also reduce reward seeking across addictive behaviors, including cocaine seeking in rodents and alcohol consumption/craving in humans^52,59^. Together, these findings suggest that GLP-1 receptor agonists may act not only by reducing appetite but by reshaping neural circuits that govern reward valuation, motivation, and behavioral termination. In this context, our identification of DMS GABAergic dynamics provides a circuit-level framework for understanding how metabolic interventions may suppress maladaptive reward-driven consumption, including overeating and potentially drug seeking, by shifting striatal output away from engagement and toward cessation.

## Acknowledgements

This work was supported by NIH grants P30DA013429 and MH132052 to MGK and T32DA007237 to BPP and the Brain and Behavior Research Foundation Young Investigator Award to MGK.

## Data availability statement

All data in the manuscript or the supplementary material are available from the corresponding author upon reasonable request. Correspondence and requests for materials should be addressed to Munir Gunes Kutlu.

## Code availability

The custom codes used for analyzing the datasets reported in this study will be provided upon reasonable request.

## Materials and Methods

### Animals

Adult (8 weeks and older) male and female C57BL/6J mice (see figure captions for number of mice for each experiment) were obtained from Jackson Laboratories (Bar Harbor, ME, SN: 000664), with 3-4 animals housed per cage, and were maintained on a 12-hour reverse light/dark cycle, with all behavioral testing occurring during the dark cycle. Animals had ad libitum access to food and water in their home cages, except during food-deprivation studies. During food deprivation, animals received only one pellet per animal per day. All experiments were conducted in accordance with the guidelines of the Institutional Animal Care and Use Committee (IACUC) at Temple University – Lewis Katz School of Medicine

### Apparatus

Fiber photometry, optogenetics, and *in vivo* calcium imaging recordings were carried out in related behavioral apparatuses, including operant conditioning chambers (Med Associates Inc., St. Albans, Vermont), a sociability chamber (60×40×22 cm, Ugo Basile SRL, Gemonio, Italy), and an open field (40×40×22 cm, Ugo Basile SRL, Gemonio, Italy). Each operant conditioning box (#ENV-307-CT, Med Associates Inc., St. Albans, Vermont) was fitted with a retractable sipper (#ENV-352AW, Med Associates Inc., St. Albans, Vermont) and a lickometer (#ENV-250, Med Associates Inc., St. Albans, Vermont).

### Surgical procedure

At least 30 minutes prior to surgery, mice were administered Meloxicam (2 mg/kg) via subcutaneous injection. Animals were anesthetized via inhaled isoflurane (3-5% for induction and 1-3% for maintenance) and placed onto a stereotaxic frame (David Kopf Instruments). Ophthalmic ointment was continuously applied to the eyes throughout the surgery to prevent corneal drying. After fur was removed from the scalp, a midline incision was made, and a craniotomy was performed with a dental drill. A 10-µL Nanofil Hamilton syringe (WPI; #NANOFIL) with a 33-gauge beveled metal needle (WPI; #NF33BV) were used to infuse viral constructs into the dorsolateral (DLS) and dorsomedial striatum (DMS). All surgeries were performed following aseptic technique.

A fluorescent γ-aminobutyric acid (GABA) sensor, pGP-AAV5-syn-iGABA.Sn.FR2-WPRE^60^ (AddGene viral prep #218874-AAV5), was unilaterally infused into the DLS (coordinates relative to bregma: anterior/poster, +0.38 mm; medial/lateral, -2.5 mm; dorsal/ventral, -3.3 mm) or DMS (coordinates relative to bregma: anterior/poster, +0.38 mm; medial/lateral, -1.5 mm; dorsal/ventral, -3.3 mm) at a rate of 50 nL/min with a total volume of 500 nL. Following viral infusion, the needle was left in the injection site for 7 minutes before slowly withdrawing.

For single-cell calcium imaging experiments, mice received unilateral infusions of AAV1.CaMK2a.GCaMP6m.WPRE.SV40 (500 nL; Addgene, catalog # if applicable) into the DMS using the same stereotaxic coordinates described above. Following viral infusion, an Inscopix ProView GRIN lens with baseplates (0.5 mm diameter, 4.0 mm length, 0.5 NA; #1050-004417, Inscopix) was implanted immediately dorsal to the injection site and secured to the skull using adhesive cement (C&B Metabond; Parkell). Mice were allowed at least 5 weeks to recover following surgery before imaging experiments commenced.

In optogenetics surgeries, an even ratio mixture of AiP11851-pAAV-hDLX-minBG-iCre-4X2C-WPRE3-BGHpA^38^ (Addgene viral prep #164450-PHPeB) and either pAAV-Syn-FLEX-rc[ChrimsonR-tdTomato]^61^ (Addgene viral prep #62723-AAV5) or pAAV-hSyn-DIO-mCherry (Addgene viral prep #50459-AAV5) was infused to the DMS at a rate of 50 nL/min with a total volume of 700 nL. For validation of optogenetic manipulations, *Chrimson* and *mCherry* mice also received a 500 nL infusion of pGP-AAV5-syn-iGABA.Sn.FR2-WPRE (AddGene; 218874-AAV5). Following this method, we were able to validate that optogenetic stimulation of GABAergic interneurons elicits reproducible, time-locked increases in GABA release across animals.

Following viral infusion, fiber-optic cannulas (Doric Lenses; 400 µm core diameter, 0.48 NA) were implanted into the DLS or DMS and positioned immediately dorsal to the viral injection site before permanently adhering to the skull using adhesive cement (C&B Metabond; Parkell). Post-operative care was performed in accordance with IACUC. Animals were allowed a minimum of 5 weeks to recover and ensure efficient viral expression before beginning experiments.

### Histology

At the end of fiber photometry, optogenetics, and calcium imaging experiments as described, animals were deeply anesthetized with isoflurane and transcardially perfused with 10 mL of 1x PBS followed by 10 mL of cold 4% PFA in 1x PBS. Animals were then quickly decapitated, brains were extracted and placed in a 4% PFA solution and stored at 4 °C for 48 hours. Brains were then transferred to a 30% sucrose solution in 1x PBS and allowed to sit until the brains sank to the bottom of the conical tube at 4 °C. After sinking, brains were sectioned at 40 μm on a freezing cryostat (Leica CM3050 S). Sections were stored in a cryoprotectant solution (7.5% sucrose + 15% ethylene glycol + 0.4% sodium azide in 0.1 M PB) at -20 °C until immunohistochemical processing.

Viral expression was validated by immunohistochemical staining. We stained all DLS and DMS slices with an anti-GFP antibody (Chicken anti-GFP; Millipore Sigma, #06-896; 1:500 in 5% bovine serum albumin [BSA]; 4 °C overnight) to validate iGABA.Sn.FR2 and GCaMP6m expression. Sections were incubated with secondary antibodies (GFP: rabbit anti-chicken CF 488A [Millipore Sigma; #SAB4600052]; 1:1000 in 5% BSA) overnight at 4 °C. Following washing, sections were counterstained for nuclei with Invitrogen^TM^ DAPI (1mg/mL at 1:6000 dilution, 5 minutes at room temperature [Fisher Scientific; #D1306]), mounted in VECTASHIELD Vibrance® Antifade Mounting Medium (#H-1700-10; Vector Laboratories; Malvern, Pennsylvania) with DAPI for nuclei counterstaining. Fluorescent images were taken using a Keyence BZ-X800 inverted fluorescence microscope (Keyence), under dry 4x and 20x objective lenses (Nikon). Viral infection and fiber optic placements were determined with serial imaging in all animals, and sections were identified that displayed the DLS or DMS, viral expression, and implant tip.

To assess the exclusion of striatal projection neurons from minBG-iCre-driven expression in our optogenetic studies, sections were immunostained for ChrimsonR-tdTomato using an anti-mCherry antibody (goat anti-mCherry; Thermo Fisher Scientific, #PA5-143590; 1:1,000 in 5% BSA) and for DARPP-32, a marker of striatal projection neurons (rabbit anti-DARPP32; Abcam; AB40801-1001; 1:500 in 5% BSA). Sections were subsequently incubated with goat anti-rabbit CF 647 (Life Technologies (Thermofisher), A32733TR) overnight at 4 °C. Imaging was performed as described above. FIJI (ImageJ) was used to quantify the total number of ChrimsonR-expressing cells and the percentage of ChrimsonR-expressing cells that colocalized with DARPP-32. Minimal colocalization with DARPP-32 was interpreted as evidence that minBG-iCre-driven expression largely excluded striatal projection neurons.

### Fiber Photometry

Viral infusion and expression of the GABA sensor iGABA.Sn.FR2 (pGP-AAV-syn-iGABA.Sn.FR2-WPRE (AddGene viral prep #218874-AAV5) in DLS and DMS neurons allowed us to record fluorescent signals through a permanently implanted fiber optic (Doric Lenses; 400 µm core diameter, 0.48 NA) secured to the skull. Our fiber photometry system utilizes two light-emitting diodes at 465nm (Tucker-Davis Technologies; #Lx465) and 405nm (Tucker-Davis Technologies; Lx405) as an isosbestic control channel, which are controlled by an LED driver (Tucker-Davis Technologies; Lux RZ10x). LED emissions pass through filter cubes and are reflected off a series of dichroic mirrors (Doric, Fluorescence MiniCube; Tucker-Davis Technologies, Lux RZ10X), allowing for emission and recording excitation through the same optical system. LEDs were controlled via real-time signal processing (Tucker-Davis Technologies, RZ10x), and emission signals from LED stimulation were determined via multiplexing. Fluorescent signals were collected via a photoreceiver (Tucker-Davis Technologies, LxPS2). To control the timing and sensitivity of LEDs as well as to record emitted fluorescence signals, Synapse software (Tucker-Davis Technologies) was utilized. The LED intensity was continuously monitored via real-time fluorescence amplitude within the Synapse software. Intensities were maintained at constant values between 350mV and 500mV across all trials for each subject. For events of interest (e.g., licks), transistor-transistor logic (TTL) signals were used to timestamp onset times from the Med-PC V software (Med Associates Inc.) to be detected via the RZ10X system in the Synapse software.

### Optogenetic Fiber Photometry

Mice were infused in the DMS with a viral mixture of AiP11851-pAAV-hDLX-minBG-iCre-4X2C-WPRE3-BGHpA (AddGene viral prep #164450-PHPeb) and pAAV-Syn-FLEX-rc[ChrimsonR-tdTomato] (AddGene viral prep #62723-AAV5) to selectively infect GABAergic interneurons (GINs) to excite GINs via optogenetic photostimulation (*Chrimson* animals). Control animals received AiP11851-pAAV-hDLX-minBG-iCre-4X2c-WPRE-BGHpA and pAAV-hSyn-DIO-mCherry (*mCherry* animals). All animals were also injected with pGP-AAV5-syn-iGABA.Sn.FR2-WPRE to allow for fiber photometry recordings of DMS GABA from optogenetically stimulated GINs. A 400µm fiber optic was also implanted into the DMS. GABA release from optogenetically excited GINs was validated (589nm, 10 sec, 10Hz, 5mW). Mice with no iGABA.Sn.FR2 signal or *Chrimson* mice with no iGABA.Sn.FR2 increases from optogenetic excitation were removed from the study.

### Single-cell calcium imaging

During each imaging session, the miniature microscope was attached to the previously implanted baseplate (see Surgical Procedures). Imaging parameters (LED power, gain, and focus) were optimized for each animal during the initial recording session and maintained constant across all subsequent imaging sessions. Data were acquired using Inscopix Data Acquisition Software (IDAS, Inscopix) while animals were freely moving alone in an operant box (MED Assoc). For single-cell calcium imaging experiments, mice received intraperitoneal injections of saline or semaglutide (0.026 mg/kg) once daily for three consecutive days, and DMS calcium activity was recorded under freely moving baseline conditions following the third treatment. At the conclusion of each recording session, the miniature microscope was removed, and the protective baseplate cover was replaced.

Calcium imaging data were acquired at 20 frames/s using an Inscopix nVoke miniature microscope and Inscopix Data Acquisition Software (IDAS, Inscopix). Image preprocessing was performed using Inscopix Data Processing Software (IDPS, Inscopix). Raw videos were spatially downsampled 2x, corrected for dropped frames, and motion corrected using a reference frame-based registration algorithm. Images were subsequently cropped to remove motion correction borders and regions outside the field of view before being exported as TIFF stacks for further analysis. Neuronal fluorescence signals were extracted using constrained non-negative matrix factorization for microendoscopic data (CNMF-E). Individual neuronal spatial footprints and calcium traces were manually inspected within the IDPS interface, and duplicated cells, neuropil contamination, and imaging artifacts were excluded from further analyses. ΔF/F traces were calculated by IDPS as the relative change in fluorescence from baseline across the imaging session. Raw CNMF-E fluorescence traces were used for all subsequent analyses.

### Behavioral Experiments

All behavioral assays were recorded using a USB camera at a top-down angle (Stoelting, #60516; TDT iVn, iV2; Inscopix, nVision).

#### Open Field Test

Animals were placed in the testing area by themselves and allowed to freely explore for 10 minutes. The average distance traveled, speed, and immobility were determined using DeepLabCut.

#### Real-Time Conditioned Place Aversion (CPA)

The biased conditioned place preference/aversion paradigm protocol was adapted from an earlier study published by our group.^62^ Training and testing were conducted in a chamber composed of two distinct compartments with distinct sensory cues (smooth floor or floor with holes) to determine whether or not optogenetic stimulation causes an aversion. Animals were individually tested and allowed to freely explore both compartments for 10 minutes. The time animals spent in each compartment was recorded to determine which side animals had a bias. The next day, animals were allowed to freely explore the test chamber and compartments. Mice received optogenetic stimulation when they entered their preferred compartment (5mW; 5 seconds on, 5 seconds off). The time spent in each compartment was timed manually by a scorer blind to the group assignments.

#### Sipper Training

Animals were trained to consume solutions of varying palatability from a retractable sipper (#ENV-352AW, Med Associates Inc., St. Albans, Vermont) in the operant conditioning chambers (#ENV-307-CT, Med Associates Inc., St. Albans, Vermont) for 30 minutes of unrestricted daily access until they reached or exceeded the minimum criterion of 500 licks. Fiber photometry animals were initially trained on 30% sucrose before other liquids of varying palatability were tested. However, 1:1 mixture of water and Ensure (Fisher Scientific; #NC1225012) was used for optogenetic animals’ initial training.

#### Sipper access with ad libitum feeding and food deprivation

After animals successfully reached or exceeded the criteria, they were exposed to various palatable solutions for 30-minutes, and animals were exposed to one type of palatable liquid per day. Animals were first given access to palatable liquids in the following order, water, 0.1% saccharin in water (Millipore-Sigma; #240931), and a 1:1 mixture of water and Ensure (Fisher Scientific; NC1225012). This testing process of different palatable liquids was first performed under an *ad libitum* diet, then repeated under food deprivation to enhance motivational states.

#### Sipper access with chronic semaglutide injections

Following a 5-day recovery period from food deprivation, during which animals had ad libitum access to food, they were tested to examine the effects of GLP-1 receptor agonism. Animals were administered subcutaneously with Semaglutide sodium (Selleck Chemicals #E7333) in a dosage of 0.026 mg/kg for 3 days. Animals were tested with 30 minutes of unrestricted access to Ensure after a 1-hour incubation period following injections. Following a 14-day washout period, the animals underwent the same testing procedure after being administered 0.9% saline solution.

#### Optogenetic Stimulation of DMS GABAergic Interneurons

Mice received optogenetic stimulations from a fiber-coupled diode laser (589nm, OptoEngine LLC, #MGL-F-589-100mW) powered by a dedicated laser power supply (Model PSU-II-LED, OptoEngine LLC). Laser stimulation timing and pulse parameters (e.g., frequency, pulse duration, and train duration) were controlled via Pulse Pal stimulator (Sanworks LLC). Laser intensity was tested daily prior to experiments using a power meter (PM100D, ThorLabs) to ensure the laser intensity was consistent across trials and experiments.

#### Open Field Test with optogenetic excitation of GABAergic interneurons

Mice were allowed to freely explore the open field for 10 minutes. As they explored, mice received 30 seconds of photostimulation (589nm, 30 sec, 10Hz, 5mW) with a 30-second interval between stimulations.

#### Conditioned Place Aversion with optogenetic excitation of GABAergic interneurons

Mice were allowed to freely explore the two compartments with different floor sensory cues in each compartment. On the first day of testing, no photostimulation was delivered. On the second day of testing, mice received 5-second long photostimulations (589nm, 5sec, 10Hz, 5mW) with a 5-second interval between stimulations when mice entered their preferred compartment.

#### Sipper access with optogenetic excitation of GABAergic interneurons

Mice were trained on the sipper as described previously. After meeting criteria, they were given one additional 30-minute sipper access session to get a baseline number of licks. The following day, mice were retested with each lick being coupled to an optogenetic stimulation (589nm, 5mW, 10Hz, 1 sec), and the number of licks was recorded. Four days later, mice were given another 30-minute sipper access session to get a baseline number of licks, then on the following day, mice were retested with optogenetic stimulations (589nm, 5mW, 10Hz, 10 sec), uncoupled and with pseudo-random intervals between laser stimulations (20, 30, or 45 seconds). All animals received a total of 50 stimulations

#### Deep Lab Cut markerless tracking analysis

Mouse locomotor activity during the open field test was recorded using an overhead camera at 15–30 frames/s. Animal position was tracked using DeepLabCut (Python 3, version 2.2b8). A ResNet-50 network was trained using >250 manually annotated frames sampled from representative videos for 200,000 iterations. Four body parts were tracked for each mouse, and only coordinates with a tracking likelihood ≥0.9 were included in the analysis.

For each body part, frame-to-frame movement was calculated as the Euclidean displacement between consecutive x–y positions:

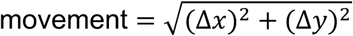

Movement values were then averaged across the four tracked body parts, ignoring missing values. This average movement trace was smoothed using a 2-frame moving window. Total distance traveled was calculated as the sum of the average frame-to-frame movement across the session and is reported in pixels. Mean speed was calculated as the average frame-to-frame movement across the session. We also calculated mean and median speed during periods of active locomotion, defined as frames in which smoothed movement was ≥5 pixels/frame. Immobility was defined as frames in which smoothed movement was <2 pixels/frame and was expressed as the percentage of total analyzed frames.

### Data Analysis and Statistics

#### Data pre-processing

Fiber photometry data analysis was conducted using a custom MATLAB pipeline as described previously^63,64^. Raw 465 nm (F465 channel) and isosbestic 405 nm (F405 channel) fluorescence signals were acquired at 1,000 Hz (1 kHz). The raw fluorescence traces from each channel were minimally smoothed using a LOWESS filter (smoothing factor = 0.0004) before ΔF/F was calculated using polynomial fitting of the 405 nm control signal to the 465 nm signal. ΔF/F was calculated as (F465 − F405)/F405. The isosbestic 405 nm channel was used to correct for GABA-independent signal fluctuations, including motion artifacts and photobleaching. Behavioral events were identified by TTL pulses generated at the onset of each lick. Peri-event ΔF/F traces were extracted from 2 s before to 10 s after each lick. The 2-s pre-lick period served as the baseline for z-score normalization, calculated as z = (signal − baseline mean)/baseline standard deviation, where the signal represents the ΔF/F value at each time point within the peri-event window. Individual peri-lick traces were then averaged within each animal to generate mean event-aligned responses for statistical analyses.

#### Peri-lick GABA signal analysis

Peri-lick GABA responses were quantified by calculating the area under the curve (AUC) from z-scored fluorescence traces using trapezoidal numerical integration over predefined post-lick time windows. AUC values were averaged across licks within each animal, and all statistical analyses were performed using the animal as the experimental unit.

#### Temporal analysis of lick GABA responses

To examine how DMS GABA dynamics evolved during ongoing consumption, lick-triggered responses were analyzed across the first 80 licks of each session, grouped into bins of 10 consecutive licks (1–10, 11–20, …, 71–80). For selected analyses, responses during the first five licks and the final lick bin (71–80) were also compared directly. To account for differences in total lick number between animals, additional analyses were performed by dividing each recording session into four equal lick quartiles.

#### Non-linear curve-fitting for GABA lick responses

To compare the temporal evolution of GABA responses across experimental conditions, the mean GABA signal across lick bins was fit using nonlinear exponential regression according to the model (y(x)=c+A(1-e^{-kx})), where (c) represents the baseline offset, (A) the asymptotic change in GABA signal, and (k) the recovery rate constant. Curves were fit separately for each condition using nonlinear least-squares regression. Differences between curves (ad libitum vs. food deprivation and saline vs. semaglutide) were assessed using an extra sum-of-squares F-test comparing a full model, in which each condition was fit with independent parameters, to a restricted model in which parameters were shared across conditions. The estimated lick bin at which the fitted curve returned to baseline (zero-crossing) was also calculated from the fitted model.

#### Statistical modeling of lick bout initiation from GABA signals

To determine whether DMS GABA dynamics predicted lick bout initiation, we used logistic regression to classify bout timing from moment-to-moment GABA signals. Whole-session ΔF/F traces were first z-scored within each recording session and downsampled from 1,000 Hz to 50 Hz. The first 2 s of each recording were removed prior to analysis. Lick bouts were defined as sequences of at least 5 licks, with individual licks separated by no more than 1 s, and bouts separated from one another by at least 10 s. Bout onset was defined as the first lick in each detected bout.

For each time point, lagged GABA features were extracted from the preceding 500 ms of the z-scored GABA trace using 20-ms steps. First-order differences of these lagged features were also included in the model. Positive labels were assigned to time points occurring within 0.75 s after bout onset, and all other time points were labeled as non-bout periods. Model performance was evaluated using leave-one-mouse-out cross-validation. In each fold, data from one mouse were held out completely as the test set, while the model was trained on data from all remaining mice. Feature standardization was performed using the mean and standard deviation calculated exclusively from the training animals and was then applied to the held-out mouse. Logistic regression models were fit using ridge regularization. To reduce class imbalance, negative samples were subsampled during training at a 5:1 ratio relative to positive samples, excluding negative samples within ±0.2 s of positive events. Negative subsampling was repeated 20 times within each cross-validation fold, and predicted probabilities were averaged across repetitions before model performance was calculated for the held-out mouse. Thus, each mouse contributed a single independent estimate of model performance.

To determine whether predictive performance reflected temporally specific information in the GABA signal, model performance was compared with two control analyses. For the permuted-label control, training labels were randomly shuffled before model fitting while the held-out mouse remained unchanged. For the time-shift control, behavioral labels were circularly shifted by 10 s within each recording session relative to the GABA signal, thereby preserving the temporal structure and boundaries of individual recordings. ROC AUC and PR AUC values were calculated separately for each held-out mouse, with mice serving as the independent unit for group-level statistical analyses.

#### Analysis of GABA signal at lick onset

To determine whether licking preferentially occurred during periods of reduced GABA signaling, we quantified the GABA dF/F value at each lick onset. For each animal, lick onset timestamps were paired with the corresponding whole-session GABA dF/F trace. The GABA signal at each lick was calculated by linearly interpolating the dF/F trace at the exact lick onset time. Licks were then classified based on whether the interpolated dF/F value was below or above zero, corresponding to below-baseline or above-baseline GABA signaling. For each animal, we calculated the number and percentage of licks occurring during below-baseline versus above-baseline GABA states. Distributions of lick-associated dF/F values were also plotted for individual animals and pooled across the cohort.

#### Detection and analysis of whole-session GABA transients

Raw ΔF/F traces were analyzed using a custom MATLAB pipeline to quantify whole-session GABA transients. The first 2 s of each recording were excluded to remove acquisition-related artifacts. Fluorescence traces were smoothed using a 50-ms moving average before baseline correction. A rolling baseline was estimated using a 45-s moving median filter, and the residual signal was calculated by subtracting this baseline from the smoothed trace. Local noise was estimated using a 45-s rolling median absolute deviation (MAD), which was converted to a local standard deviation to generate a local z-score trace. Adaptive peak detection was then performed on the local z-score signal using a minimum peak height of 2.6 standard deviations and a minimum peak separation of 100 ms. Peak widths were measured at half-height, and peak amplitudes were quantified from both the local z-score and raw ΔF/F residual signals. For each recording, the total number of whole-session GABA transients, mean and median peak amplitude, peak width, and variability of these measures were calculated and averaged within animals. In addition, inter-event intervals (IEIs), the coefficient of variation of IEIs, the fraction of short IEIs (<0.5 s), and the Fano factor of event counts (10-s bins) were calculated to quantify the temporal organization of whole-session GABA activity. The relationship between peak amplitude and duration was assessed using both Pearson and Spearman correlation coefficients as well as linear regression. To determine whether whole-session GABA dynamics changed over the recording session, the number and amplitude of detected transients were compared between the first and last 10 min of each recording (or the first and second halves for recordings shorter than 20 min). These measures were compared between mice following 3 days of saline or semaglutide treatment using the statistical tests described in the Statistical Analysis section.

#### Short vs long lick bout analysis

To examine the relationship between GABA signaling and behavioral pauses, lick-triggered responses were aligned to inter-lick intervals (ILIs). Short ILIs (<0.5 s) were classified as continuous licking, whereas long ILIs (>5 s) represented pauses in consumption. GABA responses preceding short and long ILIs were compared within animals. Finally, the probability of lick initiation occurring during periods of below-baseline GABA signaling was calculated and compared across experimental conditions.

Statistical analyses were performed using GraphPad Prism (GraphPad Software) and MATLAB. Depending on the experimental design, paired t-tests, repeated-measures ANOVA followed by appropriate multiple-comparisons tests, nonlinear regression analyses, or logistic regression were used. Statistical significance was defined as *p* < 0.05.

#### Single-cell calcium imaging and neural ensemble analysis

Custom MATLAB scripts were used for all analyses. Raw calcium traces from accepted neurons were z-score normalized across time prior to analysis. Neurons with missing values or no measurable activity were excluded. Pairwise Pearson correlation coefficients were calculated between all neuron pairs, and the mean pairwise correlation was computed from the upper triangle of the correlation matrix after excluding self-correlations. Population synchrony was quantified as the variance of the mean population activity trace across all neurons.

To characterize ensemble organization, hierarchical agglomerative clustering (average linkage) was performed on a correlation-based distance matrix (1 − Pearson’s r). Neurons with pairwise correlations ≥0.30 were grouped into the same functional ensemble. The number of ensembles, mean ensemble size, mean within-ensemble correlation, and mean between-ensemble correlation were calculated for each recording.

Population dimensionality was assessed using principal component analysis (PCA). The percentage of variance explained by the first principal component (PC1) and the first three principal components (PC1–3) was calculated. The participation ratio, defined as

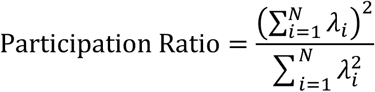

where *λ_i_*_"_is the eigenvalue corresponding to the *i*th principal component, was used as a measure of the effective dimensionality of population activity. Neural state-space volume was quantified as the square root of the determinant of the covariance matrix of the first three principal component scores. Functional connectivity networks were generated by thresholding the pairwise correlation matrix (r ≥ 0.30). Graph-theoretical analyses were performed on the resulting undirected networks to quantify graph density, mean node degree, clustering coefficient, modularity, number of connected communities, and total number of network edges. All ensemble, dimensionality, and graph metrics were compared between saline and semaglutide conditions using paired statistical analyses.

## References

1. Berthoud, H.-R. Metabolic and hedonic drives in the neural control of appetite: Who’s the boss? Curr. Opin. Neurobiol. 21, 888–896 (2011).

2. Morton, G. J., Meek, T. H. & Schwartz, M. W. Neurobiology of food intake in health and disease. Nat. Rev. Neurosci. 15, 367–378 (2014).

3. Ferrario, C. R. et al. Homeostasis Meets Motivation in the Battle to Control Food Intake. J. Neurosci. 36, 11469–11481 (2016).

4. Cox, J. & Witten, I. B. Striatal circuits for reward learning and decision-making. Nat. Rev. Neurosci. 20, 482–494 (2019).

5. Kravitz, A. V. & Kreitzer, A. C. Striatal mechanisms underlying movement, reinforcement, and punishment. Physiol. Bethesda Md 27, 167–177 (2012).

6. Balleine, B. W., Delgado, M. R. & Hikosaka, O. The Role of the Dorsal Striatum in Reward and Decision-Making. J. Neurosci. 27, 8161–8165 (2007).

7. Moyer, J. T., Halterman, B. L., Finkel, L. H. & Wolf, J. A. Lateral and feedforward inhibition suppress asynchronous activity in a large, biophysically-detailed computational model of the striatal network. Front. Comput. Neurosci. 8, 152 (2014).

8. Burke, D. A., Rotstein, H. G. & Alvarez, V. A. Striatal Local Circuitry: A New Framework for Lateral Inhibition. Neuron 96, 267–284 (2017).

9. Kramer, P. F., Twedell, E. L., Shin, J. H., Zhang, R. & Khaliq, Z. M. Axonal mechanisms mediating γ-aminobutyric acid receptor type A (GABA-A) inhibition of striatal dopamine release. eLife 9, e55729 (2020).

10. Sharott, A. et al. Different Subtypes of Striatal Neurons Are Selectively Modulated by Cortical Oscillations. J. Neurosci. 29, 4571–4585 (2009).

11. Straub, C. et al. Principles of Synaptic Organization of GABAergic Interneurons in the Striatum. Neuron 92, 84–92 (2016).

12. Gittis, A. H., Nelson, A. B., Thwin, M. T., Palop, J. J. & Kreitzer, A. C. Distinct Roles of GABAergic Interneurons in the Regulation of Striatal Output Pathways. J. Neurosci. 30, 2223–2234 (2010).

13. Mallet, N., Moine, C. L., Charpier, S. & Gonon, F. Feedforward Inhibition of Projection Neurons by Fast-Spiking GABA Interneurons in the Rat Striatum In Vivo. J. Neurosci. 25, 3857–3869 (2005).

14. Holly, E. N. et al. Striatal Low-Threshold Spiking Interneurons Regulate Goal-Directed Learning. Neuron 103, 92–101.e6 (2019).

15. Oldenburg, I. A. & Ding, J. B. Cholinergic modulation of synaptic integration and dendritic excitability in the striatum. Curr. Opin. Neurobiol. 21, 425–432 (2011).

16. Nelson, A. B. et al. Striatal Cholinergic Interneurons Drive GABA Release from Dopamine Terminals. Neuron 82, 63–70 (2014).

17. Lozovaya, N., Eftekhari, S. & Hammond, C. The early excitatory action of striatal cholinergic-GABAergic microcircuits conditions the subsequent GABA inhibitory shift. *Commun*. Biol. 6, 723 (2023).

18. Taverna, S., Ilijic, E. & Surmeier, D. J. Recurrent Collateral Connections of Striatal Medium Spiny Neurons Are Disrupted in Models of Parkinson’s Disease. J. Neurosci. 28, 5504–5512 (2008).

19. Gerfen, C. R. & Bolam, J. P. The Neuroanatomical Organization of the Basal Ganglia. in Handbook of Behavioral Neuroscience vol. 24 3–32 (Elsevier, 2016).

20. Melzer, S. et al. Distinct Corticostriatal GABAergic Neurons Modulate Striatal Output Neurons and Motor Activity. Cell Rep. 19, 1045–1055 (2017).

21. Vachez, Y. M. et al. Ventral arkypallidal neurons inhibit accumbal firing to promote reward consumption. Nat. Neurosci. 24, 379–390 (2021).

22. Giossi, C., Bahuguna, J., Rubin, J. E., Verstynen, T. & Vich, C. Arkypallidal neurons in the external globus pallidus can mediate inhibitory control by altering competition in the striatum. Proc. Natl. Acad. Sci. 121, e2408505121 (2024).

23. Frost Nylén, J., et al. The roles of surround inhibition for the intrinsic function of the striatum, analyzed in silico. Proc. Natl. Acad. Sci. 120, e2313058120 (2023).

24. Corbit, L. H. & Janak, P. H. Posterior dorsomedial striatum is critical for both selective instrumental and Pavlovian reward learning. Eur. J. Neurosci. 31, 1312–1321 (2010).

25. Balleine, B. W. & O’Doherty, J. P. Human and rodent homologies in action control: corticostriatal determinants of goal-directed and habitual action. Neuropsychopharmacol. Off. Publ. Am. Coll. Neuropsychopharmacol. 35, 48–69 (2010).

26. Yin, H. H. & Knowlton, B. J. The role of the basal ganglia in habit formation. Nat. Rev. Neurosci. 7, 464–476 (2006).

27. Kreitzer, A. C. & Malenka, R. C. Striatal plasticity and basal ganglia circuit function. Neuron 60, 543–554 (2008).

28. Berridge, K. C., Robinson, T. E. & Aldridge, J. W. Dissecting components of reward: ‘liking’, ‘wanting’, and learning. Curr. Opin. Pharmacol. 9, 65–73 (2009).

29. Villavicencio, M., Moreno, M. G., Simon, S. A. & Gutierrez, R. Encoding of Sucrose’s Palatability in the Nucleus Accumbens Shell and Its Modulation by Exteroceptive Auditory Cues. Front. Neurosci. 12, 265 (2018).

30. Joshi, A., Schott, M., la Fleur, S. E. & Barrot, M. Role of the striatal dopamine, GABA and opioid systems in mediating feeding and fat intake. Neurosci. Biobehav. Rev. 139, 104726 (2022).

31. Holst, J. J. The Physiology of Glucagon-like Peptide 1. Physiol. Rev. 87, 1409–1439 (2007).

32. Alhadeff, A. L., Rupprecht, L. E. & Hayes, M. R. GLP-1 Neurons in the Nucleus of the Solitary Tract Project Directly to the Ventral Tegmental Area and Nucleus Accumbens to Control for Food Intake. Endocrinology 153, 647–658 (2012).

33. Skibicka, K. P. The central GLP-1: implications for food and drug reward. Front. Neurosci. 7, (2013).

34. Dickson, S. L. et al. The Glucagon-Like Peptide 1 (GLP-1) Analogue, Exendin-4, Decreases the Rewarding Value of Food: A New Role for Mesolimbic GLP-1 Receptors. J. Neurosci. 32, 4812–4820 (2012).

35. Blundell, J. et al. Effects of once-weekly semaglutide on appetite, energy intake, control of eating, food preference and body weight in subjects with obesity. Diabetes Obes. Metab. 19, 1242–1251 (2017).

36. Dossat, A. M., Lilly, N., Kay, K. & Williams, D. L. Glucagon-Like Peptide 1 Receptors in Nucleus Accumbens Affect Food Intake. J. Neurosci. 31, 14453–14457 (2011).

37. Schmidt, H. D. et al. Glucagon-Like Peptide-1 Receptor Activation in the Ventral Tegmental Area Decreases the Reinforcing Efficacy of Cocaine. Neuropsychopharmacology 41, 1917–1928 (2016).

38. Graybuck, L. T. et al. Enhancer viruses for combinatorial cell-subclass-specific labeling. Neuron 109, 1449–1464.e13 (2021).

39. Cole, S., Stone, A. D. & Petrovich, G. D. The dorsomedial striatum mediates Pavlovian appetitive conditioning and food consumption. Behav. Neurosci. 131, 447–453 (2017).

40. Furlong, T. M., Jayaweera, H. K., Balleine, B. W. & Corbit, L. H. Binge-Like Consumption of a Palatable Food Accelerates Habitual Control of Behavior and Is Dependent on Activation of the Dorsolateral Striatum. J. Neurosci. 34, 5012–5022 (2014).

41. DiFeliceantonio, A. G., Mabrouk, O. S., Kennedy, R. T. & Berridge, K. C. Enkephalin Surges in Dorsal Neostriatum as a Signal to Eat. Curr. Biol. 22, 1918–1924 (2012).

42. London, T. D. et al. Coordinated Ramping of Dorsal Striatal Pathways preceding Food Approach and Consumption. J. Neurosci. 38, 3547–3558 (2018).

43. Wang, A. Y., Miura, K. & Uchida, N. The dorsomedial striatum encodes net expected return, critical for energizing performance vigor. Nat. Neurosci. 16, 639–647 (2013).

44. Yin, H. H., Ostlund, S. B., Knowlton, B. J. & Balleine, B. W. The role of the dorsomedial striatum in instrumental conditioning. Eur. J. Neurosci. 22, 513–523 (2005).

45. Spector, A. C., Klumpp, P. A. & Kaplan, J. M. Analytical issues in the evaluation of food deprivation and sucrose concentration effects on the microstructure of licking behavior in the rat. Behav. Neurosci. 112, 678–694 (1998).

46. Davis, J. D. & Perez, M. C. Food deprivation-and palatability-induced microstructural changes in ingestive behavior. Am. J. Physiol.-Regul. Integr. Comp. Physiol. 264, R97–R103 (1993).

47. Stratford, T. R. & Kelley, A. E. GABA in the Nucleus Accumbens Shell Participates in the Central Regulation of Feeding Behavior. J. Neurosci. 17, 4434–4440 (1997).

48. Pulman, K. G. T., Somerville, E. M. & Clifton, P. G. Intra-accumbens baclofen, but not muscimol, mimics the effects of food withdrawal on feeding behaviour. Pharmacol. Biochem. Behav. 97, 156–162 (2010).

49. Mietlicki-Baase, E. G. et al. Glucagon-Like Peptide-1 Receptor Activation in the Nucleus Accumbens Core Suppresses Feeding by Increasing Glutamatergic AMPA/Kainate Signaling. J. Neurosci. 34, 6985–6992 (2014).

50. Hayes, M. R. & Schmidt, H. D. GLP-1 influences food and drug reward. Curr. Opin. Behav. Sci. 9, 66–70 (2016).

51. Godschall, E. N. et al. A brain reward circuit inhibited by next-generation weight-loss drugs in mice. Nature 654, 1055–1064 (2026).

52. Hernandez, N. S. et al. GLP-1 receptor signaling in the laterodorsal tegmental nucleus attenuates cocaine seeking by activating GABAergic circuits that project to the VTA. Mol. Psychiatry 26, 4394–4408 (2021).

53. Merkel, R. et al. An endogenous GLP-1 circuit engages VTA GABA neurons to regulate mesolimbic dopamine neurons and attenuate cocaine seeking. Sci. Adv. 11, eadr5051 (2025).

54. Graham, D. L. et al. A novel mouse model of glucagon-like peptide-1 receptor expression: a look at the brain. J. Comp. Neurol. 528, 2445–2470 (2020).

55. Sandoval-Rodríguez, R. et al. D1 and D2 neurons in the nucleus accumbens enable positive and negative control over sugar intake in mice. Cell Rep. 42, (2023).

56. Shen, W., Flajolet, M., Greengard, P. & Surmeier, D. J. Dichotomous Dopaminergic Control of Striatal Synaptic Plasticity. Science 321, 848–851 (2008).

57. Barbera, G. et al. Spatially Compact Neural Clusters in the Dorsal Striatum Encode Locomotion Relevant Information. Neuron 92, 202–213 (2016).

58. Wilding, J. P. H. et al. Once-Weekly Semaglutide in Adults with Overweight or Obesity. N. Engl. J. Med. 384, 989–1002 (2021).

59. Hendershot, C. S. et al. Once-Weekly Semaglutide in Adults With Alcohol Use Disorder: A Randomized Clinical Trial. JAMA Psychiatry 82, 395–405 (2025).

60. Kolb, I. et al. iGABASnFR2 is an improved genetically encoded protein sensor of GABA. eLife 14, RP108319 (2026).

61. Klapoetke, N. C. et al. Independent optical excitation of distinct neural populations. Nat. Methods 11, 338–346 (2014).

62. Kutlu, M. G., Ortega, L. A. & Gould, T. J. Strain-Dependent Performance in Nicotine-Induced Conditioned Place Preference. Behav. Neurosci. 129, 37–41 (2015).

63. Kutlu, M. G. et al. Dopamine release in the nucleus accumbens core signals perceived saliency. Curr. Biol. 31, 4748–4761.e8 (2021).

64. Dinckol, O. et al. Dorsolateral striatal acetylcholine reorganizes neural ensembles to anticipate threat. 2025.12.10.693585 Preprint at 10.64898/2025.12.10.693585 (2025).

