## Supplementary material for "Dorsomedial striatal GABA dynamics organize palatable reward consumption and are reshaped by GLP-1 receptor agonism": Supp figures 1-10

**Extended Figures**

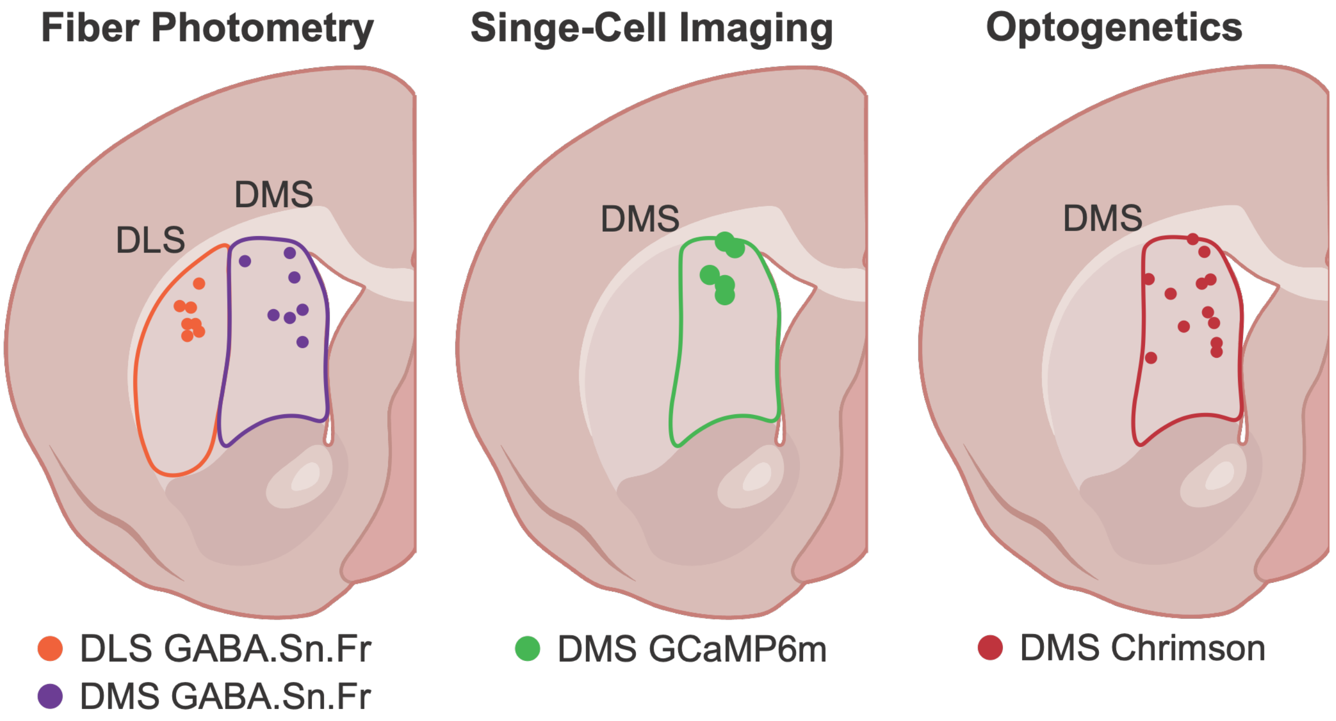


**Extended Figure 1. Placement maps for fiber photometry, single cell calcium imaging, and optogenetics experiments.** The implant placements for each experiment: **(a)** Fiber photometry implant placements in the DLS and DMS. **(b)** GRIN lens placements for the single cell calcium imaging experiments in the DMS. **(c)** Fiber optic implant placements for the DMS optogenetic experiments.

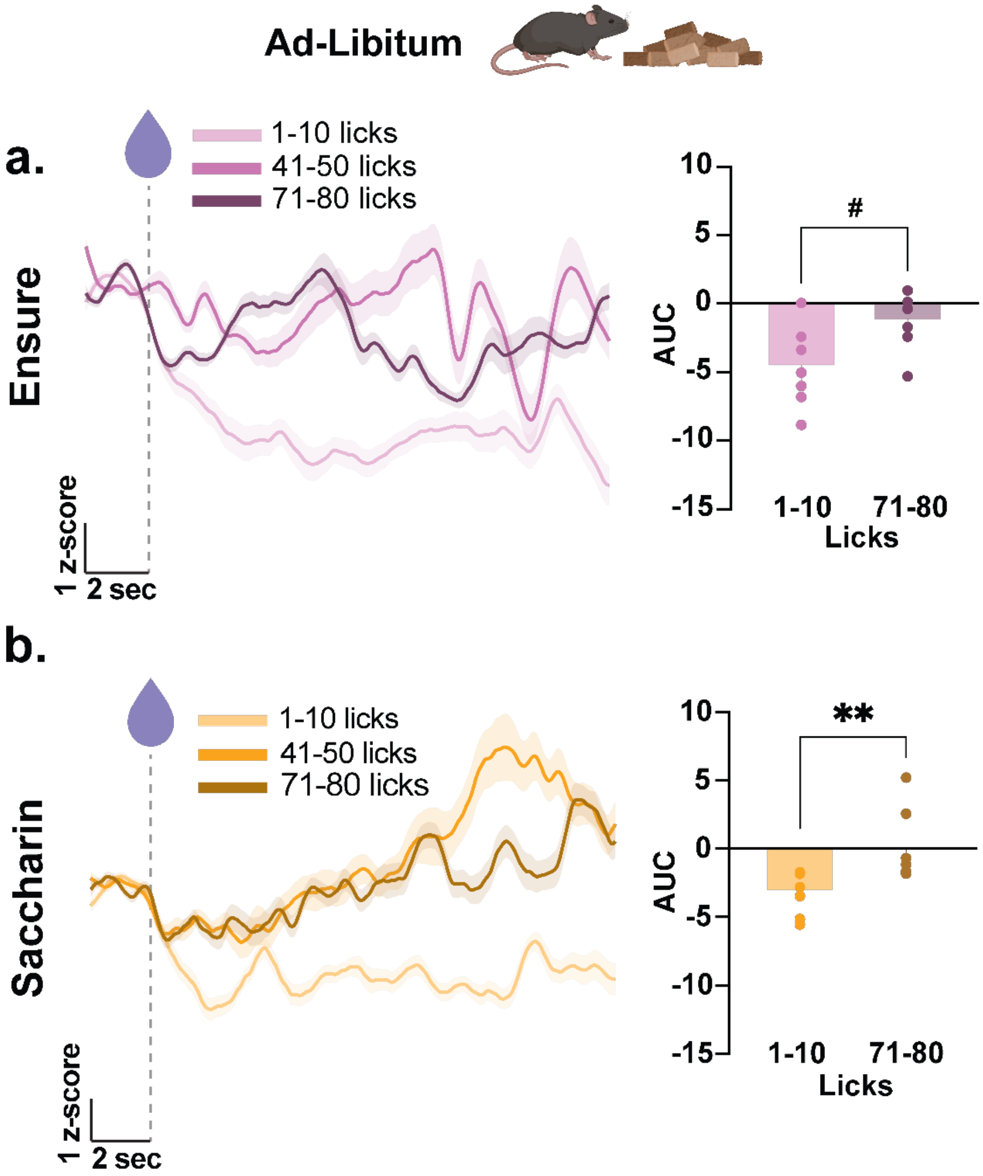


**Extended Figure 2. DMS GABA responses progressively increase across prolonged palatable liquid consumption under ad libitum feeding. (a)** Average peri-lick DMS GABA fluorescence traces during Ensure consumption under ad libitum (AL) feeding, grouped by sequential lick bins (licks 1–10, 41–50, and 71–80). Quantification of the area under the curve (AUC) comparing the first (1–10) and final (71–80) lick bins showed a trend toward increased GABA responses during late consumption (paired *t*-test, $t(6) = 2.423$, $p = 0.0516$; $n = 7$, 4 males and 3 females). **(b)** Average peri-lick DMS GABA fluorescence traces during saccharin consumption under ad libitum feeding, grouped by sequential lick bins (licks 1–10, 41–50, and 71–80). Quantification of the AUC revealed significantly greater GABA responses during the final (71–80) lick bin compared with the first (1–10) lick bin (paired *t*-test, $t(6) = 3.948$, $p = 0.0076$). The error bars represent S.E.M.s. $^{\#}p=0.0516$, $^{**}p<0.01$.

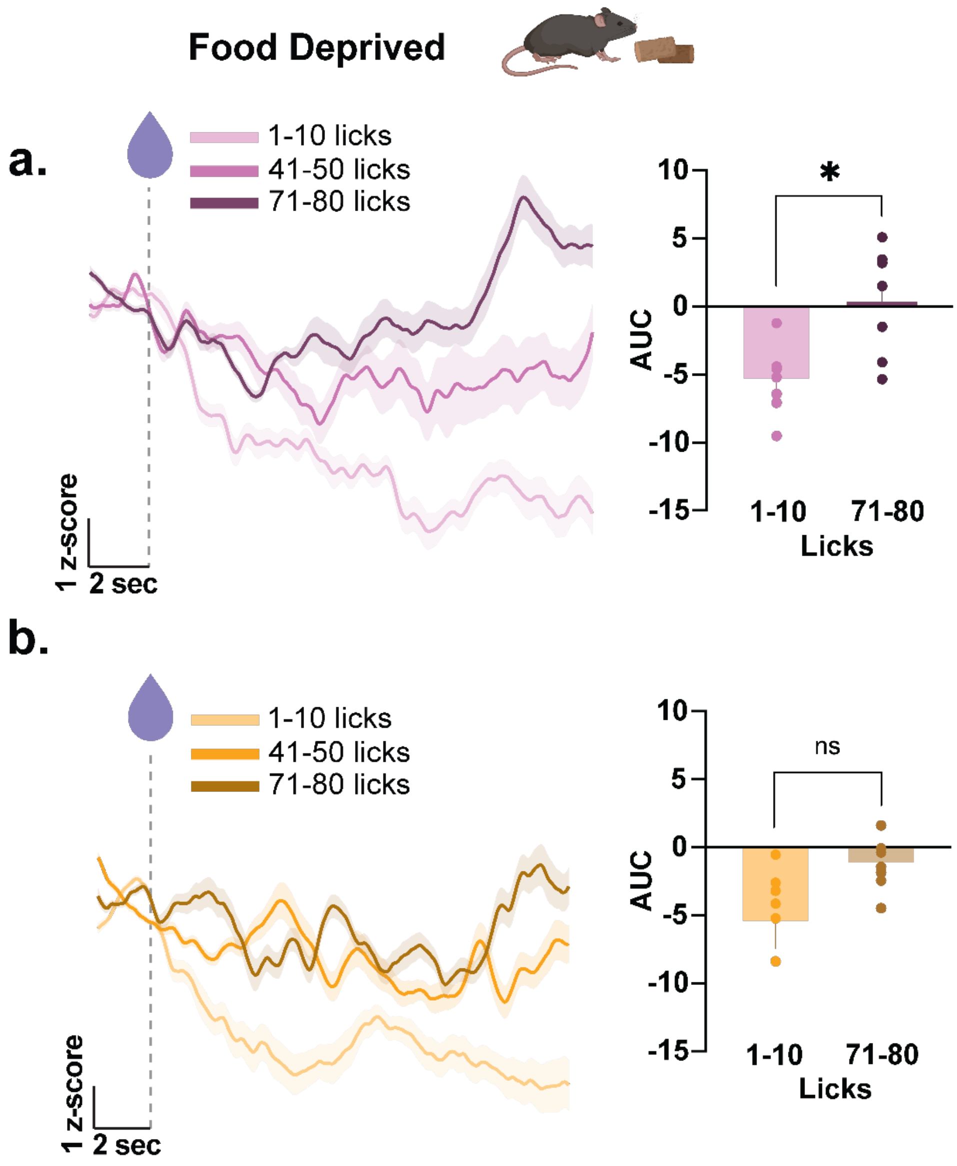


**Extended Figure 3. Food deprivation enhances late DMS GABA responses during prolonged palatable consumption. (a)** Average peri-lick DMS GABA fluorescence traces during Ensure consumption under food-deprived (FD) conditions, grouped by sequential lick bins (licks 1–10, 41–50, and 71–80). Quantification of the area under the curve (AUC) comparing the first (1–10) and final (71–80) lick bins revealed significantly greater GABA responses during late consumption (paired *t*-test, $t(6) = 2.980$, $p = 0.0246$); $n = 7$, 4 males and 3 females. **(b)** Average peri-lick DMS GABA fluorescence traces during saccharin consumption under food-deprived conditions, grouped by sequential lick bins (licks 1–10, 41–50, and 71–80). Quantification of the AUC showed no significant difference between the first (1–10) and final (71–80) lick bins (paired *t*-test, $t(6) = 1.833$, $p = 0.1165$). The error bars represent S.E.M.s. ns=not significant, $p>0.05$, $^*p<0.05$.

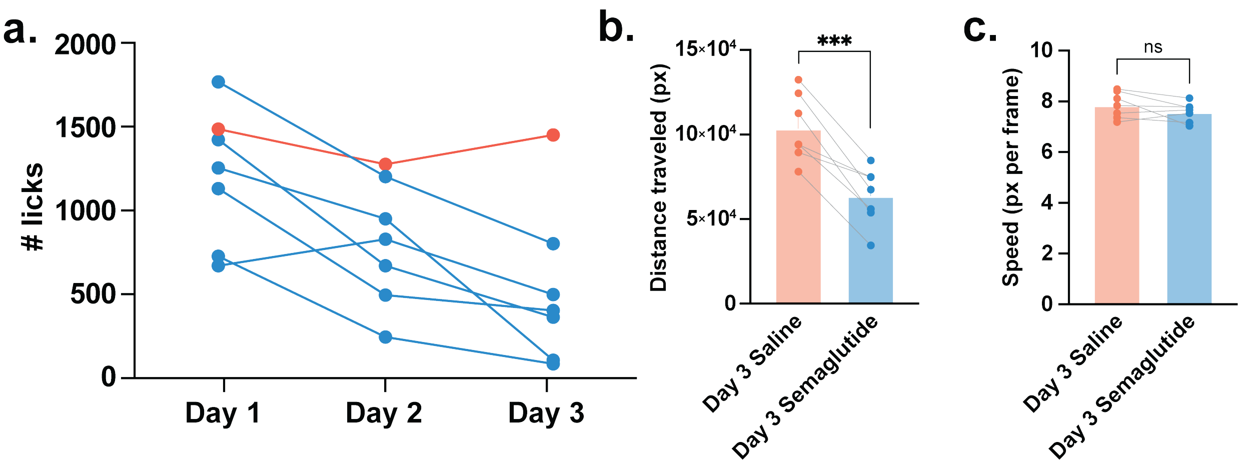


**Extended Figure 4. Chronic semaglutide reduces locomotor activity without affecting movement speed during Ensure consumption. (a)** Individual lick counts across three consecutive days of semaglutide treatment for each mouse ($n$ = 6, 3 males and 3 females), illustrating progressive reductions in Ensure consumption over the treatment period. **(b)** Total distance traveled during the 30-minute Ensure access session on Day 3 following saline or semaglutide treatment ($n$ = 7, 4 males and 3 females). Semaglutide significantly reduced locomotor activity compared with saline (paired $t$-test, $t(6) = 6.084$, $p = 0.0009$). **(c)** Mean locomotor speed during the Day 3 Ensure access session following saline or semaglutide treatment. Mean speed did not differ between treatment conditions (paired $t$-test, $t(6) = 1.462$, $p = 0.1940$). The error bars represent S.E.M.s. ns=not significant, $p>0.05$, ***$p<0.001$.

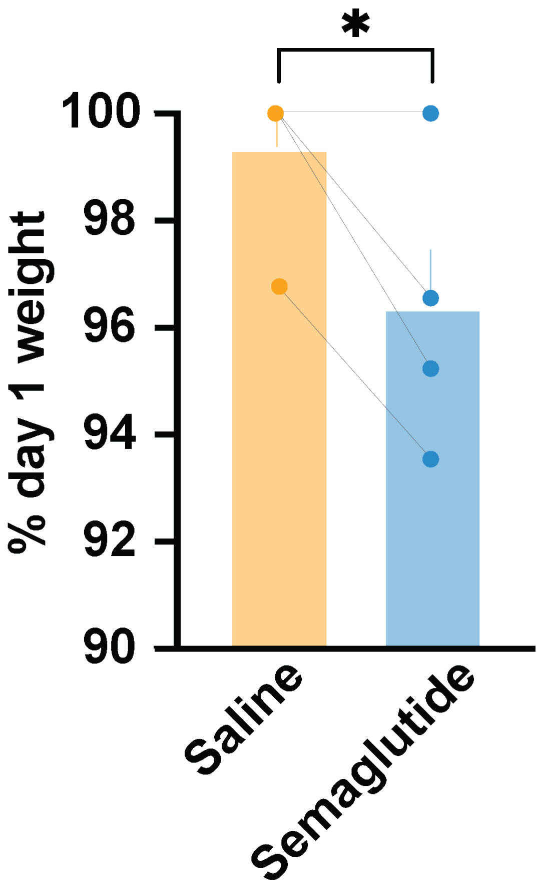


**Extended Figure 5. Chronic semaglutide reduces body weight in mice used for single-cell calcium imaging.** Body weight on Day 3 expressed as a percentage of Day 1 body weight following saline and semaglutide treatment in mice used for single-cell calcium imaging (*n* = 5, 3 males and 2 females). Semaglutide significantly reduced body weight compared with saline (paired *t*-test, $t(4) = 3.757$, $p = 0.0198$). The error bars represent S.E.M.s. ns=not significant, p>0.05, ***p<0.001.

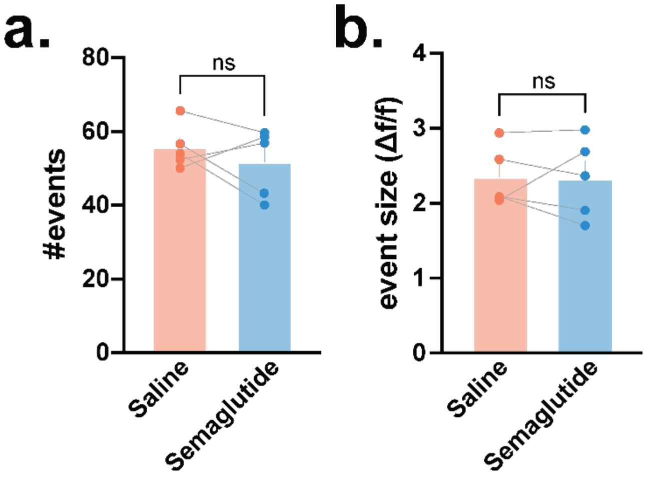


**Extended Figure 6. Quantification of calcium event frequency and event size in the DMS neural ensembles following semaglutide administration.** Semaglutide did not significantly alter either **(a)** event frequency (paired *t*-test, $t(4) = 0.8928$, $p = 0.4224$) or **(b)** event size (paired *t*-test, $t(4) = 0.1026$, $p = 0.9232$). The error bars represent S.E.M.s. ns=not significant, $p>0.05$.

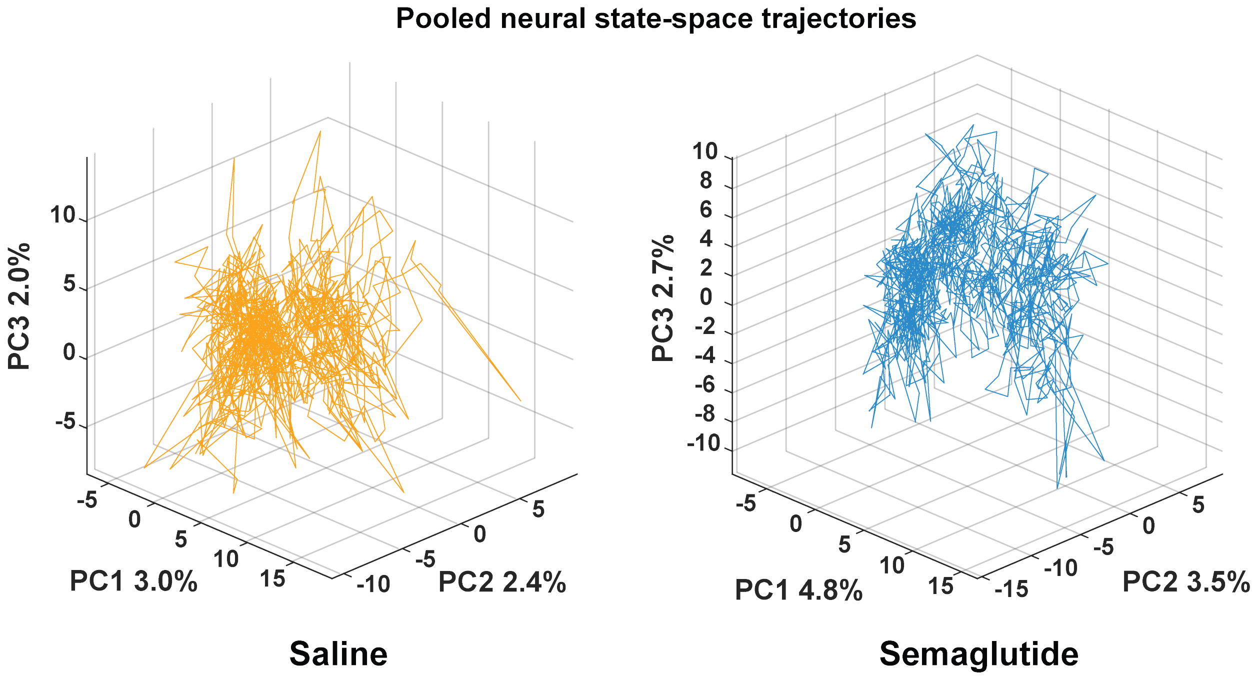


**Extended Figure 7. Neural state-space trajectories of DMS population activity following saline and semaglutide administration.** Three-dimensional principal component analysis (PCA) trajectories of pooled DMS neuronal population activity under saline (left) and semaglutide (right) treatment. Population activity from all recorded neurons was projected onto the first three principal components, illustrating the evolution of DMS network activity throughout the recording session.

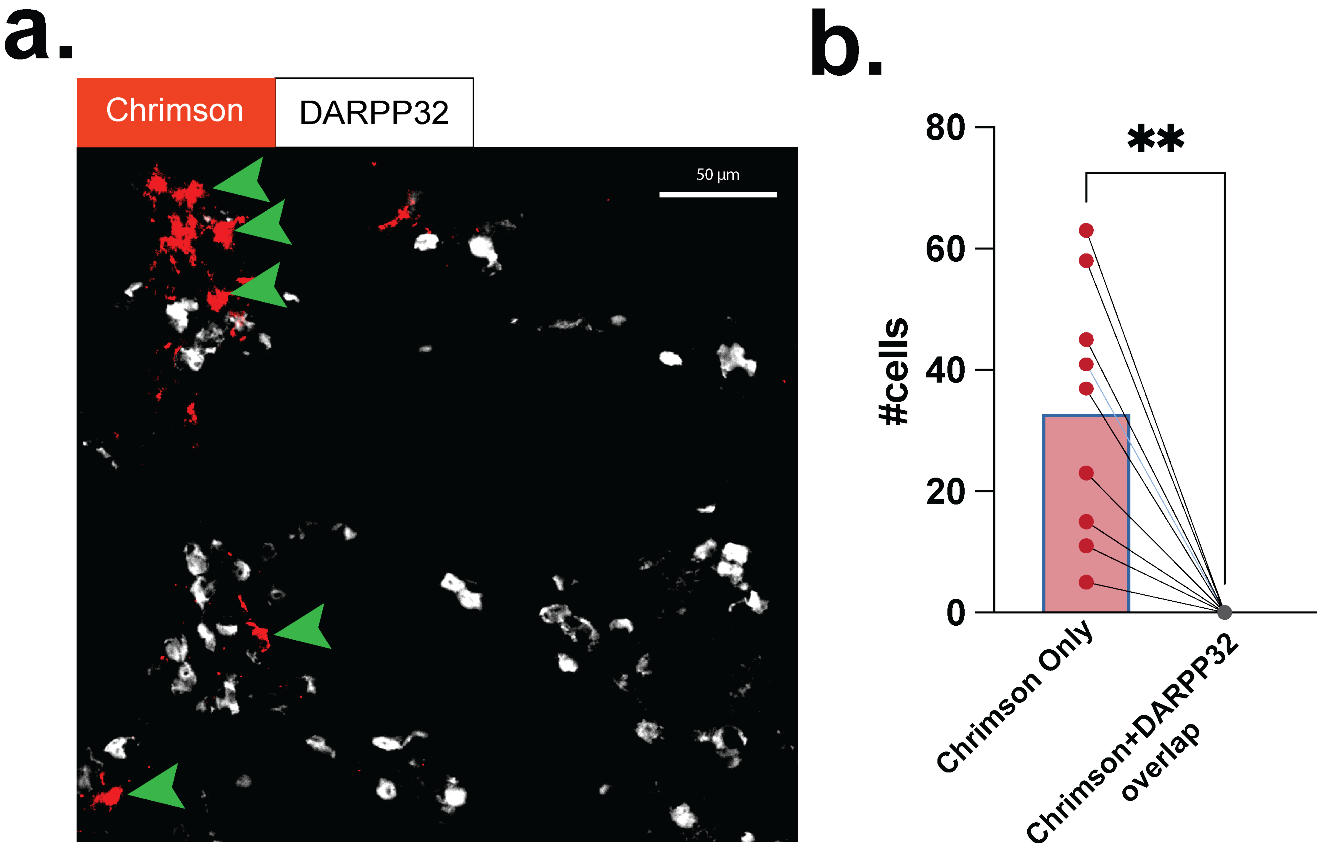


**Extended Figure 8. Histological validation of hDLX expression in the dorsal striatum. (a)** Representative fluorescence image showing hDLX (Crimson; red) expression in the dorsal striatum. Sections were immunolabeled with DARPP-32 (white), a marker of medium spiny neurons. **(b)** Quantification of hDLX-expressing (Crimson$^+$) cells and their colocalization with DARPP-32. Nearly none of the hDLX-expressing cells colocalized with DARPP-32, indicating minimal overlap between hDLX expression and DARPP-32-positive neurons. Data were analyzed using a paired two-tailed *t*-test ($t(8) = 4.790$, $p = 0.0014$; $n = 9$ brain sections). Individual data points represent biological replicates, with lines connecting paired measurements. **$p<0.01$.

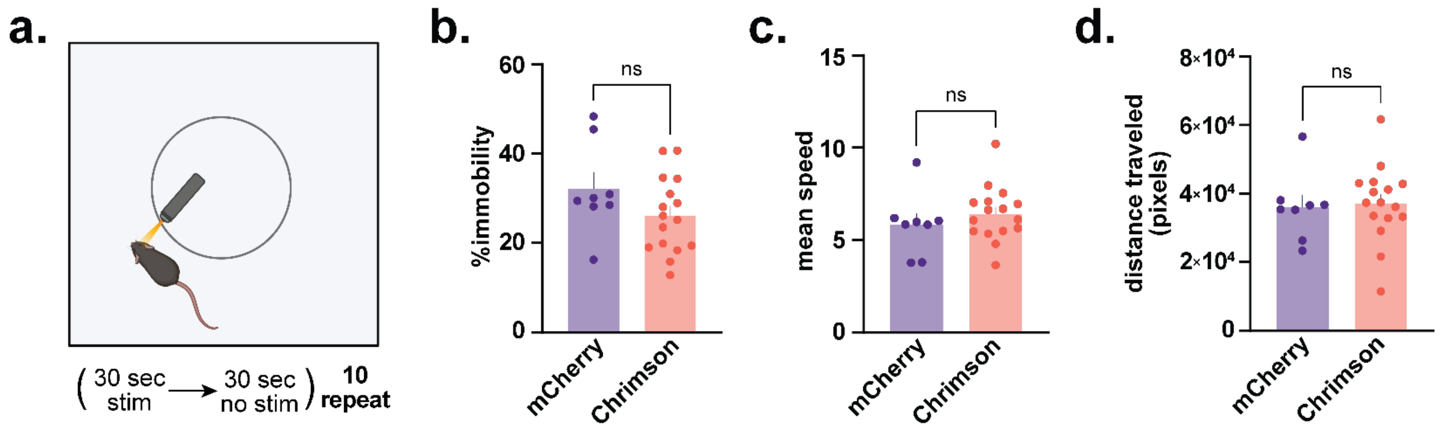


**Extended Figure 9. Optogenetic activation of DMS GABAergic interneurons does not alter locomotion or immobility in the open field. (a)** Open-field behavioral paradigm. Mice received alternating 30-s epochs of 589-nm optogenetic stimulation (5 mW, 10 Hz) and 30-s epochs without stimulation throughout the recording session to assess locomotor behavior independent of reward consumption. **(b)** Percentage of time spent immobile during the open-field session. Immobility did not differ between Chrimson (*n* = 15, 9 males and 6 females) and mCherry (*n* = 8, 4 males and 4 females) mice (unpaired *t*-test, $t(22) = 0.7471$, $p = 0.4629$). **(c)** Mean locomotor speed during the open-field session. Optogenetic activation of DMS GABAergic interneurons did not significantly alter movement speed (unpaired *t*-test, $t(22) = 0.2301$, $p = 0.8202$). **(d)** Total distance traveled during the open-field session. Total locomotor activity was comparable between Chrimson and mCherry mice (unpaired *t*-test, $t(22) = 0.7701$, $p = 0.4494$). The error bars represent S.E.M.s. ns=not significant, p>0.05.

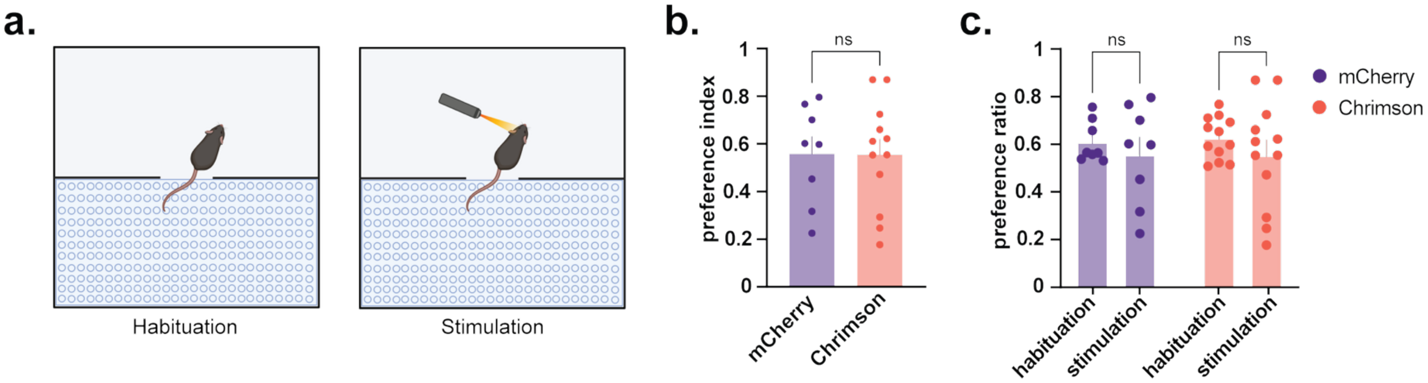


**Extended Figure 10. Optogenetic activation of DMS GABAergic interneurons does not produce conditioned place preference or aversion. (a)** Conditioned place aversion (CPA) paradigm. Mice first underwent a habituation session to assess baseline chamber preference. During the conditioning session, one chamber was paired with 589-nm optogenetic stimulation (5 mW, 10 Hz) while the opposite chamber remained unstimulated. **(b)** Preference index measured following conditioning. Preference index did not differ between Chrimson ($n$ = 12, 9 males and 3 females) and mCherry ($n$ = 8, 4 males and 4 females) mice (unpaired $t$-test, $t(18) = 0.029$, $p = 0.9771$). **(c)** Preference ratio during habituation and conditioning sessions for mCherry and Chrimson mice. Repeated-measures ANOVA revealed no significant session × genotype interaction ($F(1,18) = 0.066$, $p = 0.8000$) and no significant main effect of genotype ($F(1,18) = 0.011$, $p = 0.9172$). Sidak's multiple-comparisons tests were not significant ($p > 0.05$). The error bars represent S.E.M.s. ns=not significant, p>0.05.
